# Dissecting 16p11.2 hemi-deletion to study sex-specific striatal phenotypes of neurodevelopmental disorders

**DOI:** 10.1101/2023.02.09.527866

**Authors:** Jaekyoon Kim, Yann Vanrobaeys, Zeru Peterson, Benjamin Kelvington, Marie E. Gaine, Thomas Nickl-Jockschat, Ted Abel

## Abstract

Neurodevelopmental disorders (NDDs) are polygenic in nature and copy number variants (CNVs) are ideal candidates to study the nature of this polygenic risk. The disruption of striatal circuits is considered a central mechanism in NDDs. The 16p11.2 hemi-deletion (16p11.2 del) is one of the most common CNVs associated with NDD, and 16p11.2 del/+ mice show sex-specific striatum-related behavioral phenotypes. However, the critical genes among the 27 genes in the 16p11.2 region that underlie these phenotypes remain unknown. Previously, we applied a novel strategy to identify candidate genes associated with the sex-specific phenotypes of 16p11.2 del/+ mice and identified 3 genes of particular importance within the deleted region: thousand and one amino acid protein kinase 2 (*Taok2*), seizure-related 6 homolog-like 2 (*Sez6l2*), and major vault protein (*Mvp*). Using the CRISPR/Cas9 technique, we generated 3 gene hemi-deletion (3g del/+) mice carrying null mutations in*Taok2, Sez6l2*, and *Mvp*. We assessed striatum-dependent phenotypes of these 3g del/+ mice in behavioral, molecular, and imaging studies. Hemi-deletion of *Taok2, Sez6l2*, and *Mvp* induces sex-specific behavioral alterations in striatum-dependent behavioral tasks, specifically male-specific hyperactivity and impaired motivation for reward seeking, resembling behavioral phenotypes of 16p11.2 del/+ mice. Moreover, RNAseq analysis revealed that 3g del/+ mice exhibit gene expression changes in the striatum similar to 16p11.2 del/+ mice, but only in males. Pathway analysis identified ribosomal dysfunction and translation dysregulation as molecular mechanisms underlying male-specific, striatum-dependent behavioral alterations. Together, the mutation of 3 genes within the 16p11.2 region phenocopies striatal sex-specific phenotypes of 16p11.2 del/+ mice, unlike single gene mutation studies. These results support the importance of a polygenic approach to study NDDs and our novel strategy to identify genes of interest using gene expression patterns in brain regions, such as the striatum, which are impacted in these disorders.

## INTRODUCTION

Neurodevelopmental disorders (NDDs) are highly heritable polygenic disorders ^1-3^. Although various studies have identified genetic variants associated with NDDs, it is still unclear how sets of genes interact with each other to impact the function of distinct neuronal circuits, ultimately resulting in behavioral phenotypes. Copy number variants (CNVs) are ideal candidates to define polygenic mechanisms underlying NDDs. CNVs show high effect sizes and include multiple genes ^4-6^, allowing for the investigation of how multiple genes act together to cause vulnerability for NDD phenotypes. Although it is possible to study a single genetic variant at a time, currently there is no established strategy to identify and examine the effects of a group of candidate genes on distinct phenotypes.

16p11.2 hemi-deletion (16p11.2 del), one of the most common CNVs associated with NDDs ^7, 8^, mediates several phenotypes, including reward processing and motor activity via alterations in striatal circuits ^8-13^. Striatal dysfunction, in turn, has been observed in patients across various NDDs such as autism spectrum disorders (ASD) and attention deficit hyperactivity disorder (ADHD) ^14-25^. In humans, NDDs exhibit a male preponderance with males outnumbering females by 2 to 4 fold ^26-29^. Consistent with this, female sex was a protective factor for 16p11.2 del carriers ^13^. Of note, mice modeling 16p11.2 hemi-deletion (16p11.2 del/+ mice) show several sex-specific phenotypes ^9, 11, 30^, including male-specific deficits in behaviors heavily dependent on striatal circuits, such as reward processing and motivation ^10^. Male-specific alterations in peristriatal fiber tracts found in 16p11.2 del/+ mice could be a neuroanatomical correlate of these behavioral changes ^31^. Thus, 16p11.2 del/+ mice provide a promising model to study male-specific striatal vulnerability.

These findings indicate that mice modeling 16p11.2 del are robust translational models for studying the role of male-specific vulnerability to striatal circuits in NDDs, but little is known about the genetic mechanisms underlying the sex-specific male vulnerability or female resilience of 16p11.2 del. One particularly puzzling riddle is which of the 27 genes in the deletion contribute to these distinct sex-specific phenotypes. Single gene studies have replicated partial phenotypes of the 16p11.2 del phenotypes ^32-37^, however, these approaches are unable to examine synergistic combinatorial effects of multiple genes in this locus. Recent work underscores the idea that multiple genes within the 16p11.2 region may exhibit significant polygenic influences more broadly in ASD, thus establishing the 16p11.2 region as a source of both common and rare genetic variation ^38^. One major obstacle for the identification of synergistic oligo- or polygenic effects for a given CNV is the lack of established strategies to identify and select candidate genes. Here, we build on the findings of a previous study from our lab that used a novel strategy to identify candidate genes based on their spatial expression patterns ^31^. We selected a set of genes with expression patterns that overlapped with male-specific structural changes in striatal circuits in 16p11.2 del/+ mice to determine if haploinsufficiency of this set of genes might drive the striatum-related sex-specific behavioral phenotypes of 16p11.2 del/+ mice. These genes are thousand and one amino acid protein kinase 2 (*Taok2*), seizure-related 6 homolog-like 2 (*Sez6l2*), and major vault protein (*Mvp*). Previous studies have indicated the relevance of each of these three genes for neuronal development. *Taok2* has been associated with spine maturation and the stabilization of post-synaptic densities ^33, 34, 39^, and *Sez6l2* has been implicated in the regulation of neurite outgrowth ^40, 41^. *Mvp* is highly expressed in developing neurons, especially the dendrites of cortical neurons ^42, 43^. However, sex-specific phenotypes have not been reported in mice carrying deletions of any of these single candidate genes ^33, 35, 44^.

Considering the polygenic nature of NDDs, we hypothesized that the combination of mutations in these three candidate genes would give rise to the sex-specific phenotypes observed in 16p11.2 del/+ mice. Mice carrying mutations in all three genes cannot be produced by crossing existing single gene KO mice because of the close proximity of these genes in the genome. Therefore, we used the CRISPR/Cas9 system to develop a novel mouse model with the simultaneous hemi-deletion of these 3 candidate genes to investigate whether the combinatorial effect of these genes mediates sex-specific striatal behavioral phenotypes and transcriptomic alterations. We investigated the impact of the combinatorial effect of these genes on behavior, gene expression, and brain structure. We found that the 3 gene mutations led to sex-specific changes in behavior, specifically hyperactivity and reduced motivation for reward in males. We also found that the mutations led to similar changes in gene expression in the striatum as 16p11.2 del/+ mice in males and analysis indicates that ribosomal dysfunction could be responsible for these behavior alterations. Overall, these results highlight the value of a polygenic approach in the study of NDDs and offers fresh insights into the dissection of CNVs to identify genes of interest.

## METHODS AND MATERIALS

### Animals

All procedures were approved by the University of Iowa Institutional Animal Care and Use Committees (IACUC), and we followed policies set forth by the National Institutes of Health Guide for the care and use of laboratory animals. Mice with a selective hemi-deletion (3g del/+ mice) were generated at the Genome Editing Core of the Carver College of Medicine at University of Iowa using CRISPR/Cas9-induced gene modifications. 16p11.2 del/+ mice were purchased from The Jackson Lab (#013128). Male and female littermate animals (10-14-week-old) during the same time period were used in all experiments unless otherwise specified.

Detailed methods and materials involving animals, behavioral tests, MRI imaging, biochemical experiments, and analyses are provided in the supplement document.

### Generation of 3g del/+ mice

C57BL/6J male mice were bred with super-ovulated females to produce zygotes for electroporation. Female ICR mice were used as recipients for embryo transfer. Cas9 RNPs and the electroporation mix were prepared. Pronuclear-stage embryos were collected using methods previously described ^45^. Electroporation was performed and embryos were immediately implanted into pseudo-pregnant ICR females.

### Behavioral tasks

#### Activity monitoring

Mice locomotor activity was measured by activity monitoring using an infrared beam-break system (Opto M3, Columbus Instruments, Columbus, OH) as previously described ^9, 46^.

#### Operant task

Mice were allowed to acclimate to a 0900–2100 hours reversed light cycle one week prior to obtain food-restricted weights of 85-90% of free-feeding weights. Chocolate flavored Ensure Original Nutrition Shake (Abbott) diluted to 50% with water was used as a reinforcer. Fixed ratio (FR) responses per day for each animal were recorded and analyzed using a repeated measures ANOVA with genotype as the between-subject factors and day as the within-subjects factor, as previously described ^10^, using GraphPad 9. For progressive ratio (PR), average breakpoints were assessed by unpaired t-tests using GraphPad 9.

### RNA quantification in striatal samples

RNA isolation was conducted as previously described ^10^. The striatum was dissected bilaterally using a mouse brain matrix allowing 1 mm sections. The tissue was stored in RNAlater, then RNA was extracted using Qiagen RNeasy kit (Qiagen) according to the manufacturer’s instructions. cDNA was synthesized and Quantitative real time polymerase chain reaction (qPCR) was performed. The specific RNA expression was calculated using GraphPad Prism 9 (San Diego, CA, USA).

### RNA-seq

#### Library preparation

RNA quality was assessed and samples with RNA integrity number (RIN) >8 were used to perform RNA-seq. RNA library preparation from WT mice (n□=□8 samples, 4 males and 4 females) and 3g del/+ mice (n□=□8 samples, 4 males and 4 females) for 3g del/+ mice and WT mice (n□=□8 samples, 4 males and 4 females) and 16p11.2 del/+ mice (n□=□8 samples, 4 males and 4 females) for 16p11.2 del/+ mice were prepared using the Illumina TruSeq Stranded Total RNA with Ribo-Zero gold sample preparation kit (Illumina, Inc., San Diego, CA). Pooled libraries were sequenced on Illumina NovaSeq 6000 sequencers with 100-bp Paired-End chemistry (Illumina). The dataset is available in the NCBI’s Gene Expression Omnibus repository, GEO Series accession GSE224750.

#### RNA-seq analysis

Sequencing data was processed with the bcbio-nextgen pipeline (https://github.com/bcbio/bcbio-nextgen). All further analyses were performed using R (version 4.1.2). Analysis code available through Github at https://github.com/YannVRB/16p11.2-del-and-3gKO-bulk-RNA-seq.git

### Acquisition and analysis of diffusion-weighed MRI data sets

#### Image acquisition

Male and female mice at an age of PD 42-47 (6-week-old), 70-day-old (10-week-old), or 114-118-day-old (16-week-old) were imaged using a GE/Agilent Discovery 901 7-Telsa pre-clinical scanner. MRI imaging acquisition consisted of a 32-direction diffusion tensor imaging (DTI) scan.

#### Image preprocessing and TBSS analysis

Diffusion-weighed images were converted from DICOM to NIFTI format using DCM2NIIX and controlled for quality. We generated fractional anisotropy (FA) data set. Voxel-wise statistical analysis was performed on the FA data using a version of Tract-Based Spatial Statistics (TBSS), as previously described ^31^. Briefly, mean FA images were created and aligned FA data was projected onto the mean FA skeleton. Voxel-wise cross-subject statistics (threshold-free cluster enhancement (TFCE) at p < 0.05) were performed.

## RESULTS

### Generation of 3 gene hemi-deletion mice

We generated mice with a simultaneous hemi-mutation in each of the 3 candidate genes (*Taok2, Sez6l2*, and *Mvp*) using CRISPR/Cas9-induced gene modifications to introduce a premature stop codon in the coding region of each gene (Figure 1A and B). The targeting modification was designed to activate nonsense-mediated mRNA decay (NMD) for specific degradation of transcripts of each of the genes. For *Taok2*, a 10 bp deletion and inversion of part of the 9 bp deleted sequence were introduced in exon 2, creating a premature stop codon in exon 4 (Figure 1C and Figure S1A). We introduced a 17 bp deletion in exon 2 of *Sez6l2* to generate a frame shift mutation causing a premature stop codon in exon 3 (Figure 1D and Figure S1B). We made a 1 bp insertion in exon 2 of the *Mvp* gene to generate a premature stop codon in exon 2 (Figure 1E and Figure S1C). Sanger DNA sequencing was performed to confirm the modification in several brain regions (cortex, cerebellum, and striatum) and peripheral tissues (ear and tails). The results verified a successful modification of the candidate genes. To validate the impact of these modifications on expression levels, we analyzed mRNA levels of the target genes (*Taok2, Sez6l2*, and *Mvp*) and nearby genes (*Hirip3, Tmem219, Asphd1, Cdipt*, and *Prrt2*) in the striatum. These results confirmed that the gene modifications decreased mRNA levels of each of the 3 target genes, but did not alter the expression of nearby genes (Figure 1F-H and Figure S2). No significant fatalities of 3g del/+ pups were found (Figure S3A). Interestingly, we found a reduction in body weight in male 3g del/+ mice compared to sex- and age-matched wildtype (wt) mice at the age of 7 weeks (Figure S3B), resembling 16p11.2 del/+ male mice that show decreased body weight compared to wt mice ^11, 47, 48^. However, the weight change was not found in female 3g del/+ mice (Figure S3B), unlike 16p11.2 del/+ female mice that have lower body weight than wt female mice ^11^.

**Figure 1.**
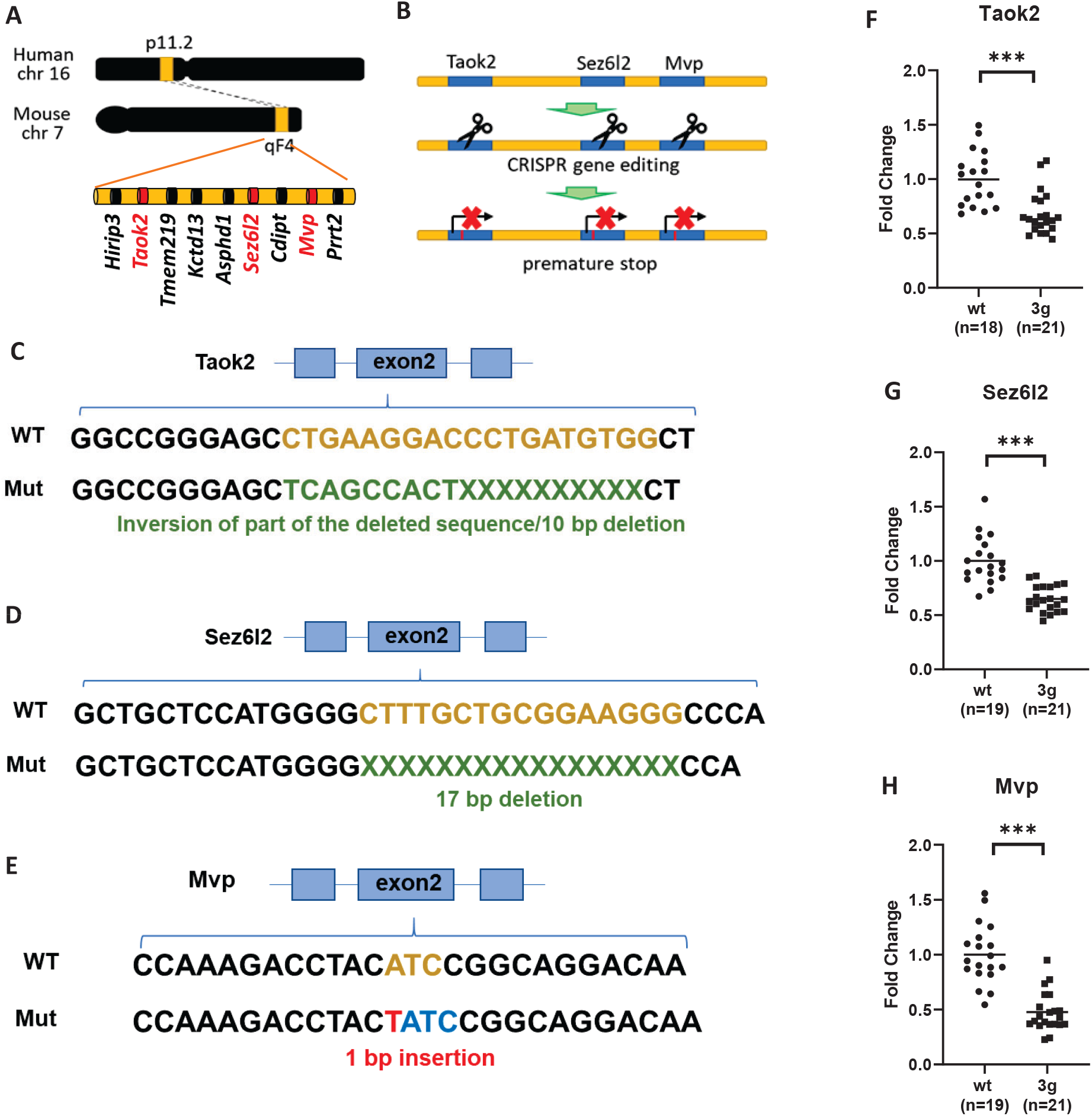
Generation of 3g del/+ mice. (A) Schematic diagram shows the target sites of mutations, mouse chromosome 7qF4. (B) A schematic presentation of the generation of 3 gene hemi-deletion mice by CRIPSR/Cas9. (C-E) Schematic presentations indicate the strategies of modifications. (F-H) qPCR results demonstrated decreased gene expressions of the 3 genes in the mutant mice (two-tailed Student t test, *** p < 0.001).

### Male-specific striatum-dependent behavioral alteration in 3 gene hemi-deletion mice

We first focused on behaviors mediated by striatal networks in the 3g del/+ mouse model, as the three candidate genes were chosen based on the overlap between their expression patterns and structural alterations in striatal circuits of 16p11.2 del/+ mice ^31^. In particular, we wanted to investigate whether 3g del/+ mice would exhibit hyperactive behavior and deficits in reward learning and motivation as observed in 16p11.2 del/+ mice ^9, 10^. First, we examined locomotor behavior in 3g del/+ mice using activity monitoring in the home-cage across the 24-hour cycle with an infrared beam-break system, as in previous experiments with 16p11.2 del/+ mice ^9^. Interestingly, male 3g del/+ mice displayed increased locomotor activity (Figure 2A), but female mice did not (Figure 2B). When activity was analyzed across the light/dark cycle, we found significantly increased activity in the dark (active) phase (Figure 2C) for male 3g del/+ mice, but not female 3g del/+ mice (Figure 2D). Although female 3g del/+ mice show a trend of hyperactive behavior with a significant behavioral difference in a few time slots, they did not fully recapitulate the hyperactivity observed in 16p11.2 del/+ females ^9^. These results point to the significance of our 3 candidate genes in the regulation of locomotor behavior regulation in male mice, phenocopying hyperactive behaviors of 16p11.2 del/+ mice. In addition, motor coordination and balance of 3g del/+ mice was evaluated in the rotarod test. 3g del/+ mice showed similar performance to wt mice in both sexes (Figure S4), suggesting that hyperactivity in male 3g del/+ mice is not linked to alterations in motor performance, coordination, or learning.

**Figure 2.**
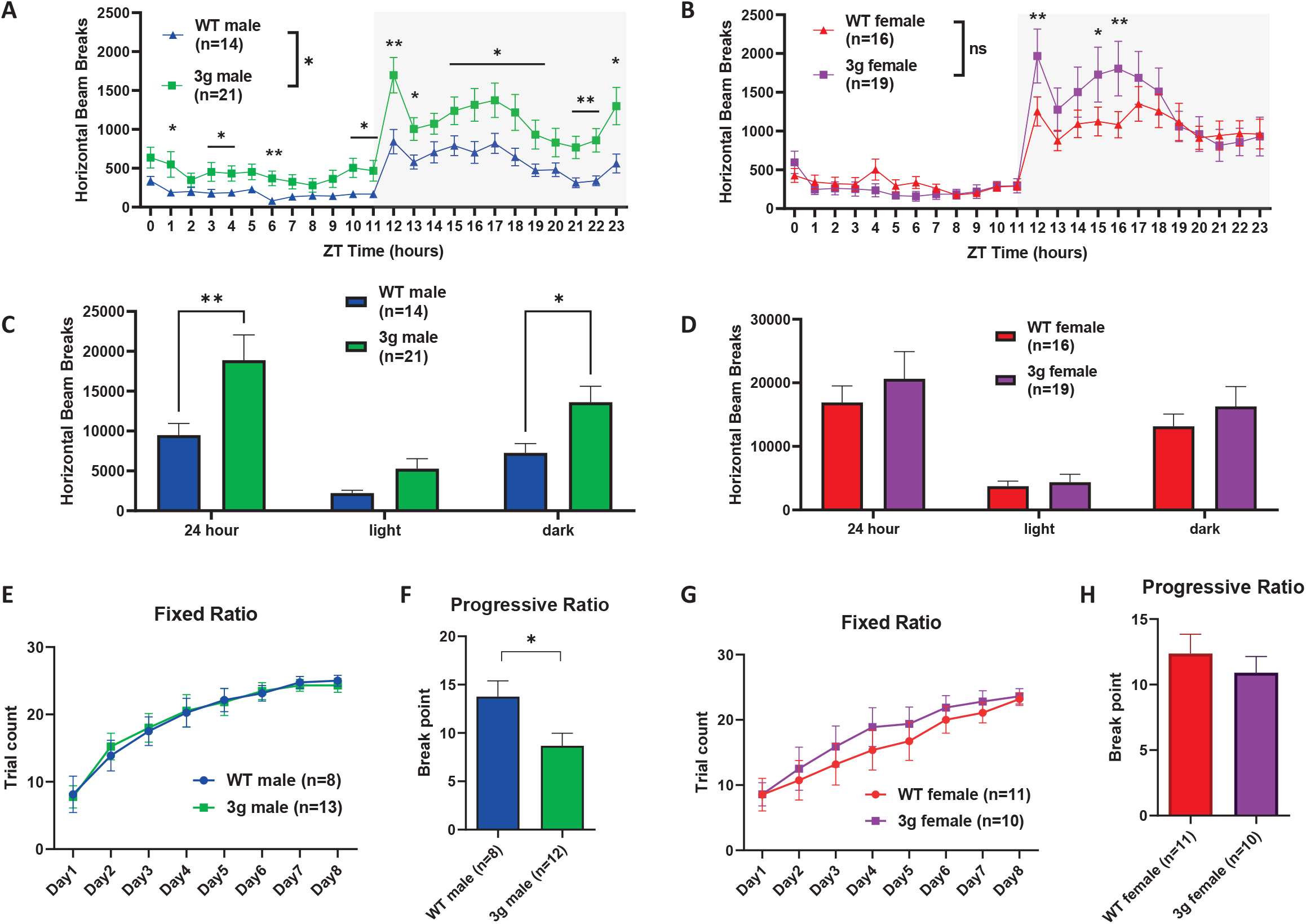
Striatum dependent behavioral alterations in 3g del/+ mice. (A, B) Infrared beam breaks in the horizontal axis are plotted for male and female 3g del/+ mice and wt mice across the 24-hr day with 1-hr bin. Gray box indicates the dark cycle. Male 3g mice have significantly greater activity relative to wt littermates (main effect of genotype, F (1, 33) = 5.373, p = 0.0268; main effect of time, F (4.368, 144.2) = 29.41, p < 0.001; genotype x time interaction, F (23, 759) = 2.249, p < 0.001), but not female 3g del/+ mice (main effect of genotype, F (1, 33) = 0.2537, p = 0.6178; main effect of time, F (23, 759) = 36.19, p < 0.001; genotype x time interaction, F (23, 759) = 2.791, p < 0.001). (C, D) Infrared beam breaks are plotted by light/dark cycle. Male 3g mice show hyperactive behavior relative to wt littermates in dark cycle (main effect of genotype, F (1, 99) = 13.60, p = 0.0004; post hoc, p = 0.034). Female 3g del/+ mice have no differences in horizontal activity (main effect of genotype, F (1, 99) = 1.256, p = 0.2650). (E, G) Both male and female 3g del/+ mice acquire a nosepoke response under a fixed ratio schedule of reinforcement indistinguishable from wt mice (male, main effect of genotype, F (1, 19) = 0.002, p = 0.968; female, main effect of genotype, F (1, 19) = 0.3530, p = 0.559). (F, H) In a progressive ratio test of motivation, male 3g del/+ mice responses after significantly fewer trials than wt males (t(18) = 2.43, p = 0.026), while female 3g del/+ mice show no different responds than wt mice (t(19) = 0.750, p = 0.463). Values are displayed as mean (±SEM) and significance values are set at *p<0.05 and **p<0.01.

Next, we investigated 3g del/+ mice using the operant reward learning task. After mild food restriction, 3g del/+ and wt mice were trained on a fixed-ratio task (FR) in mouse operant chambers. The 3g del/+ mice and wt mice showed similar weight changes after food restriction regardless of sex (Figure S5). In the FR task, 3g del/+ mice did not show differences in reward learning compared to wt mice for both sexes (Figure 2E and G), suggesting that the learning of reward-stimuli association is unaltered. A week after FR training, we tested motivation to obtain a reward using a progressive ratio schedule (PR). Interestingly, male 3g del/+ mice showed a decreased break point for the PR compared to male wt mice (Figure 2F), suggesting diminished motivation to work for reinforcement. However, female 3g del/+ mice showed no difference compared to wt females (Figure 2H). These results reveal that only male 3g del/+ mice exhibit reduced motivation to work for reinforcement. This finding is consistent with 16p11.2 del/+ mice ^10^, which show a male-specific impairment in motivation. In sum, the data from our behavioral characterization suggest that our selective 3 gene hemi-deletion phenocopies distinct male-specific behavioral phenotypes that critically depend on striatal circuits. We examined other behavioral phenotypes of the 3g del/+ mice including the open field task and social approach task, revealing male-specific reduced anxiety-like behavior and normal sociability of 3g del/+ mice (Figure S6).

### Transcriptomic changes in 3 gene hemi-deletion mice in the striatum

Patterns of gene expression are essential in mediating complex behaviors, including cognition, affective processing, and addiction ^49-51^. Identifying transcriptomic changes in the striatum of 3g del/+ mice may help us understand the mechanisms underlying the behavioral phenotypes. To determine the transcriptomic consequences of our selective 3g hemi-deletion, we compared gene expression between male and female 3g del/+ and wt mice using RNA sequencing. Our analysis revealed 817 genes differentially expressed between 3g del/+ and wt males in the striatum, and 346 genes differentially expressed between 3g del/+ and wt females in the striatum (Figure 3A, Table S1 and S2). Only 36 differentially expressed genes (DEGs) were shared between male and female 3g del/+ mice (Figure 3A, Table S5), suggesting that distinct transcriptomic changes in the striatum between male and female 3g del/+ mice may underlie the sex-specific alterations in striatal behavior.

**Figure 3.**
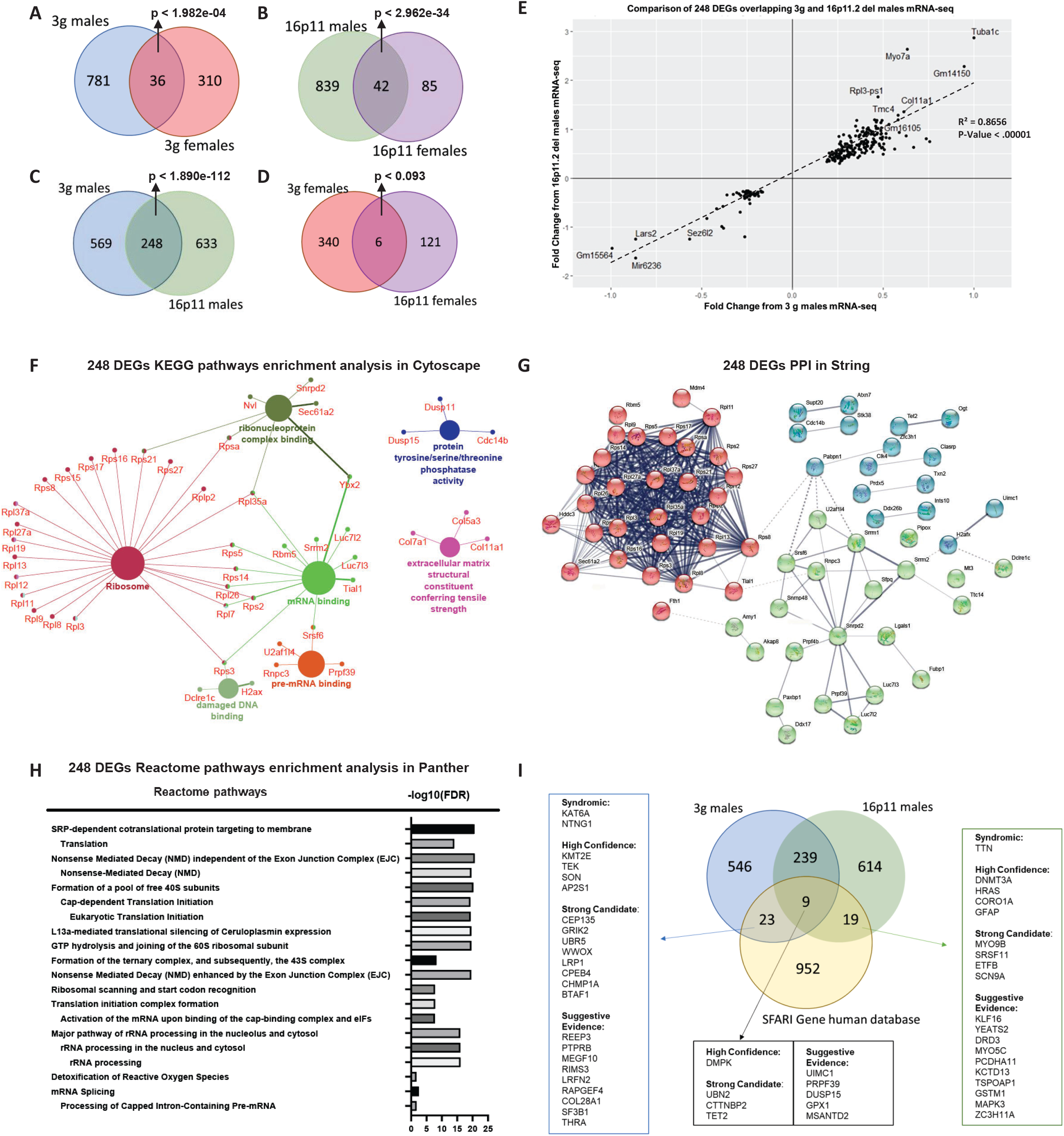
Gene expression changes in the striatum of 3g del/+ mice. (A, B) Venn diagram showing the number of overlapping DEGs that were altered in 3g del/+ mice (A) and 16p11.2 del/+ mice (B) (males and females separately). (C, D) Venn diagram representing the number of overlapping DEGs between 3g del/+ mice and 16p11.2 del/+ mice sex-specifically. (E) Quadrant plot based on comparison of genes regulated in 3g del/+ male mice and 16p11.2 del/+ male mice. Linear regression was applied that generated a coefficient of correlation which was used to calculate the p-value using Pearson (R) Calculator. The value of coefficient of determination (R²) is 0.8656 with significant p value (p < 0.00001) in regression (F) Pathway analysis using ClueGo (KEGG and GO: Molecular function terms) of 248 DEGs, the overlapping DEGs between male 3g del/+ mice with male 16p11.2 del/+ mice, shows functional groupings of network of enriched categories for the DEGs. Significant pathways are presented with a corrected p-value <0.05 (Bonferroni step down). (G) Protein-protein interaction (PPI) networks of 248 DEGs show closely connected proteins. The network is composed of 68 nodes and with a minimum required interaction score of 0.400. The nodes were drawn in three different colors dividing the interacting network into 3 clusters composed of closely connected proteins in the network. Edges of the network indicated the strength of data support. (H) An overrepresentation test on 248 DEGs was performed in PANTHER using the reactome pathway annotation dataset. Only pathways with FDR < 0.05 were displayed. (I) Venn diagram showing the overlap of significantly DEGs from 3g male del/+ mice and 16p11.2 del/+ male mice with the SFARI Gene autism susceptibility database focusing on the human gene module. The overlapping genes are categorized by SFARI respective score. The syndromic category contains genes that are linked with significantly increased risk and further characteristics not required for an ASD diagnosis. The high confidence category includes mutations that are clearly implicated in ASD by the presence of at least three de novo likely-gene-disrupting mutations being reported in the literature. Genes with two or single de novo likely-gene-disrupting mutations are categorized as “Strong candidate genes” or “Suggestive evidence”, respectively.

Next, we compared the transcriptomic signatures of our 3g hemi-deletion to the16p11.2 deletion. We performed RNA-seq from the striatum of 16p11.2 del/+ mice and littermate controls. We found 881 DEGs between male 16p11.2 del/+ mice, and 127 DEGs in 16p11.2 females with 42 overlapping DEGs, both compared to littermate controls (Figure 3B, Table S3, S4, and S6). We compared DEGs between 3g del/+ mice and 16p11.2 del/+ mice for each sex. Remarkably, there are 248 overlapping DEGs in the striatum of male 3g del/+ mice and male 16p11.2 del/+ mice (Figure 3C, Table S7, p < 1.890e-112), but only 6 overlapping DEGs between female groups (Figure 3D, Table S8, p < 0.093), supporting the hypothesis that the hemi-deletion of our 3 candidate genes plays an important role in male-specific striatal transcriptomic alterations and behavioral phenotypes observed in 16p11.2 del. Moreover, the quadrant plot of 248 DEGs indicates that all DEGs in common between 3g del/+ and 16p11.2 del/+ males change in the same direction without a single exception (Figure 3E). These results strongly support the hypothesis that the 3 gene hemi-deletion largely shares its transcriptomic fingerprint with 16p11.2 deletion in the striatum of male mice. Next, we performed a KEGG enrichment pathway analysis to identify the molecular functions of the 248 overlapping male DEGs (Figure 3F) to provide insights into potential mechanisms underlying male-specific, striatum-dependent behavioral alterations. The pathway analysis of these 248 overlapping DEGs revealed significant dysregulation of DEGs within ribosome-related pathways, with a total of 24 ribosomal genes downregulated in the male striatum of both 3g del/+ and 16p11.2 del/+ mice (Figure S7). Also, protein-protein interactions of the 248 DEGs were analyzed by a protein-protein interaction (PPI) network generated using the STRING database (Figure 3G). This analysis also implicated genes coding for ribosomal proteins involved in the cellular process of translation. Further, an overrepresentation test in PANTHER displayed significant alterations in a translation-related pathway (FDR < 6.73E-17) in the striatum of male 3g del/+ and 16p11.2 del/+ mice (Figure 3H). Together, our results indicate that changes of gene expression in ribosomal pathways linked to protein translation constitute the transcriptomic basis for striatal male vulnerability/female resilience to NDDs.

We also investigated whether DEGs in the male 3g del/+ and 16p11.2 del/+ mice showed any overlap with a database of candidate genes relevant to ASD using the SFARI Gene database. We found that 32 DEGs in 3g del/+ male mice overlapped with ASD risk genes (Figure 3I, p < 9.838e-05), and 28 overlapping DEGs in 16p11.2 del/+ male mice overlapped with ASD risk genes (Figure 3I, p < 3.608e-07). There were 9 genes that overlapped within all three categories: *DMPK, UBN2, CTTNBP2, TET2, UIMC1, PRPF39, DUSP15, GPX1, MSANTD2* (Figure 3I, p < 0.00285).

Additional pathway analyses for DEGs for 3g del/+ males only (569 DEGs, Figure S8), 16p11.2 del/+ males only (633 DEGs, Figure S9), 3g del/+ females only (340 DEGs, Figure S10), and 16p11.2 del/+ females only (121 DEGs, Figure S11) highlight the enrichment of RNA binding in 3g del/+ males and collagen modification in 16p11.2 del/+ males. We also performed pathway analysis for overlapping DEGs from Figure 3A (36 DEGs, Table S5), Figure 3B (42 DEGs, Table S6), and Figure 3D (6 DEGs, Table S8), however, they did not show significant results because of the small number of DEGs.

### Brain structural changes in 3 gene hemi-deletion mice

Previously we have shown sex-specific fiber tract changes in 16p11.2 del/+ mice using high-resolution MRI with diffusion tensor imaging (DTI) ^31^. DTI scans allow inference on the fiber tract architecture of the brain ^52^. Homologous changes in increased fractional anisotropy (FA) in medial white matter regions was reported in human 16p11.2 hemi-deletion patients ^53^. Therefore, we investigated brain structural changes of 6-week-old 3g del/+ mice using DTI as described previously ^31^.

To our surprise, we did not find any significant differences in FA between 3g del/+ mice and wt mice in male mice (Figure 4A and S12A), in contrast to previous findings in 16p11.2 del/+ mice ^31^. Only female 3g del/+ mice showed significant FA decreases in a small region of striatal fiber tracts compared to wt female mice (Figure 4B and S12B). Because our previous 16p11.2 del/+ MRI study used older mice ^31^, we subjected 6-week-old 16p11.2 del/+ mice to an identical DTI sequence to explore for age-related effects. Strikingly, in contrast to older mice, there was no significant FA difference between 16p11.2 del/+ male mice and wt mice (Figure 4C and S12C) at 6 weeks of age. However, we found widespread decreased FA in 16p11.2 del/+ females compared to wt females (Figure 4D and S12D), consistent with previous results in 10-week-old female mice ^31^. As no FA change was detected in both 6-week-old male 3g mice and 16p11.2 del mice, we inspected 16-week-old male and female 3g mice and 10-week-old male 16p11.2 del mice to examine the age-dependent brain structural changes. Consistent with our previous study, FA changes are detected in male 10-week-old 16p11.2 del mice (Figure S12G), however, FA changes are not detected in either male and female 16-week-old 3g mice at this age (Figure 4E,F and S12E,F). These findings suggest complex disruptions in the brain maturation process in 16p11.2 del/+ and 3g del/+ mice that will require further investigation.

**Figure 4.**
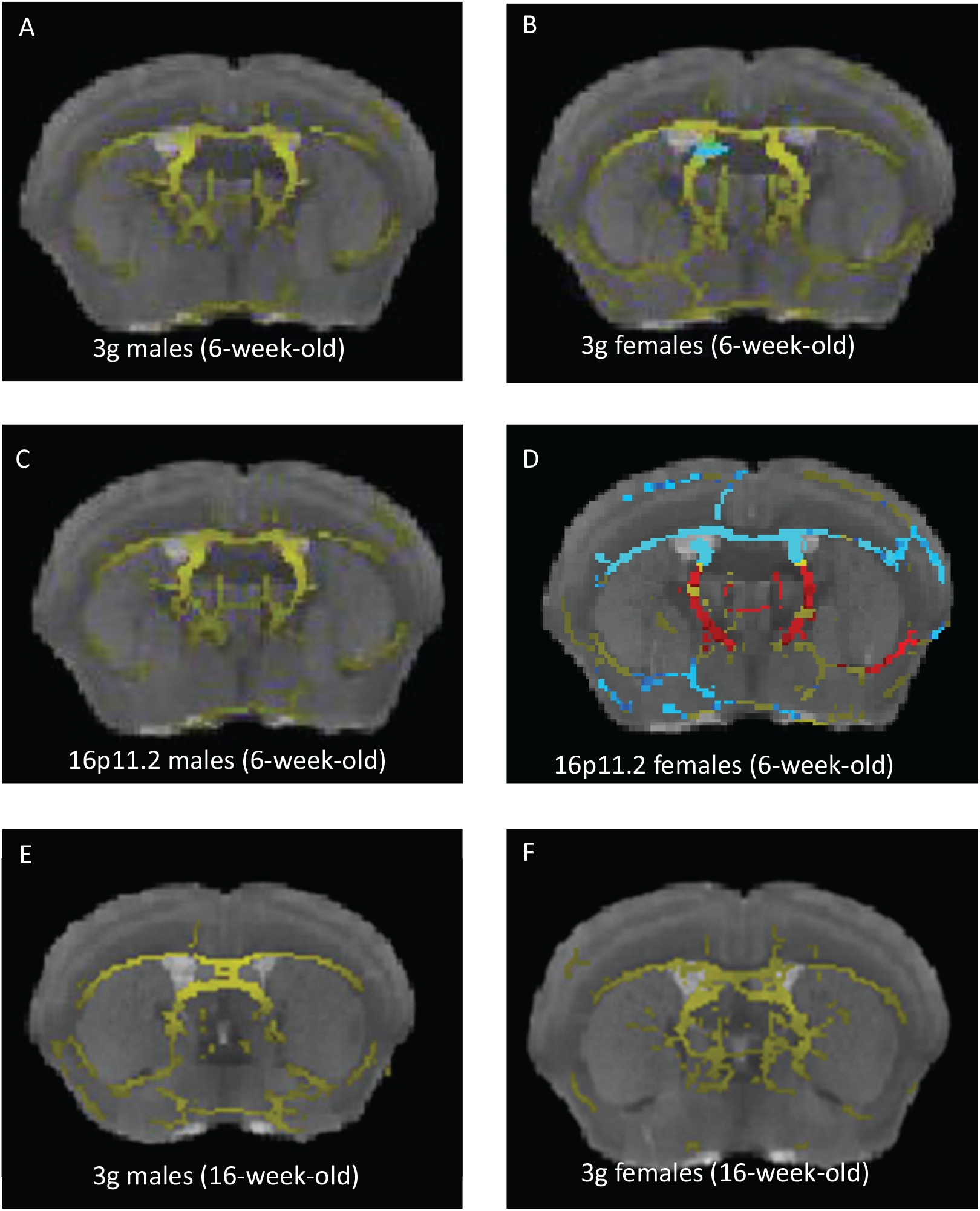
Fiber tract changes in 3g del/+ mice compared to wild types. (A-D) Increased fractional anisotropy (FA) in 3g del/+ or 16p11.2 del/+ mice compared to wt mice is represented in blue and decreased FA is displayed in red. The mean fiber tract skeleton is shown in yellow. FA alterations are not detected in 6-week-old 3g del/+ and 16p11.2 del/+ male mice (A, C), while increased FA in small region in 3g del/+ female mice (B). Widespread increased FA and pronounced FA decreases are displayed 16p11.2 del/+ female mice (D). FA alterations are not detected in 16-week-old 3g del/+ mice (E, F). Significant changes on the brain region are indicated on representative images (p<0.05).

## DISCUSSION

Disruptions of striatal circuits are found in patients with ASD ^14, 18, 19^ and ADHD ^20^, as well as genetic mouse models for the study of these disorders ^54-57^, and are related to core NDD pathologies, such as dysfunctional reward processing and altered locomotor activity. Of note, a male bias is observed in both human patients with these disorders and some of the respective mouse models. The molecular basis of male-specific striatal vulnerability, however, has not been identified.

Based on a novel strategy to identify genes-of-interest based on their spatial gene expression patterns, we identified 3 candidate genes from the 16p11.2 region, *Taok2, Sez6l2*, and *Mvp*. ^31^. Our hypothesis was that these 3 genes alone are sufficient to mediate sex-specific behavioral phenotypes that depend on striatal circuits. Our newly generated 3g del/+ mouse model displays sex-specific striatal behavioral and transcriptomic characteristics, resembling the phenotypes of 16p11.2 del/+ mice. These results demonstrate that our novel approach of selecting genes of interest based on their spatial gene expression patterns in structurally altered brain regions is a valid path to develop data-driven hypotheses that can be later tested by generating model organisms with a selective modification of the respective genes, providing novel insights into polygenic interactions underlying neurodevelopmental disorders.

Our behavioral studies reveal that a selective hemi-deletion of our 3 candidate genes leads to aspects of the striatal behavioral phenotypes also seen specifically in 16p11.2 del male mice. Previously, we found that male, but not female, 16p11.2 del/+ mice show impairments in reward-directed learning as well as motivation to work for rewards. Although 3g del/+ male mice show impaired motivation, we did not find an impairment in reward learning in 3g del/+ male mice. There are two potential explanations for the observed difference between the 3g del/+ mice and 16p11.2 del/+ mice. First, it could be because of a difference in FR protocol difficulty in the two studies. Previously, we applied the five-choice serial reaction time task (5-CSRTT) to 16p11.2 del/+ mice, a task uses 5 holes for correct 5-CSRTT performance ^10^. In this study, we tried to minimize the number of days in training and restricted access to one hole, reducing extraneous responding to non-active holes. Second, it is possible that the three genes are not sufficient to produce the entire phenotype. For instance, it was reported that a heterozygous deletion of KCTD13, a gene within 16p11.2, induced object recognition memory deficits ^32^, suggesting potential contribution of other genes related to the phenotypes of the 16p11.2 del/+ mice.

To identify the mechanisms underlying these behavioral phenotypes, we investigated transcriptomic changes in the striatum of 3g del/+ mice and 16p11.2 del/+ mice. The comparisons of striatal DEGs between male 3g del/+ mice and 16p11.2 del/+ mice revealed 248 overlapping DEGs. It is noteworthy that the overlapping DEGs between male 3g and 16p11.2 del/+ mice are statistically significantly enriched above chance (p < 1.890e-112), however, the overlapping DEGs identified in female 3g and 16p11.2 del/+ mice are not (p < 0.093). The pathway analysis of these 248 DEGs revealed that a ribosome-related pathway and translation regulation are significantly dysregulated, highlighting potential mechanisms that underlie the male-specific striatum-dependent behavioral alterations. A recent tissue-specific transcriptome profiling study on 16p11.2 del/+ mouse model also reported significant GO-term enrichments related to translation, in the striatum and cerebellum ^58^. Although they did not provide a sex specific analysis, these results support possible disruption of translational regulation in 16p11.2 del/+ mice. Reduced basal protein synthesis in the hippocampus of 16p11.2 del/+ mice were reported ^59^. The downregulation of ribosomal genes could affect translation efficiency leading to protein synthesis dysregulation and corresponding to presentation of NDDs ^60-62^. Recent studies of NDDs like Fragile X Syndrome and Tuberous Sclerosis have shown sex-specific differences in ribosomal function or translation regulation that may contribute to the sex-specific phenotypes ^63-66^. Further studies on translational regulation will provide insights into the sex-specific pathological mechanisms of NDD.

Another impactful result of the KEGG enrichment analysis for 248 DEGs is tyrosine/serine/threonine phosphatase activity related genes including *Dusp15* (Figure 3F). *Dusp15* plays an important role in regulation of extracellular signal-regulated kinase (ERK) activation ^67^. The link between *Dusp15* and ERK leads us to the idea of a potential link between 3 genes and ERK pathway. Interestingly, our 3 candidate genes are all related to the regulation of ERK signaling. MVP has a well-established association with ERK signaling, serving as a scaffold protein for ERK ^68, 69^. Taok2 has been shown to interact with Septin7 at the postsynaptic density, which can activate ERK signaling ^70^, and Sez6l2 phosphorylates PKC ^40^, which is a known activator of ERK. Although our current study does not provide a clear mechanistic explanation of the combinational effects, one of our hypotheses is the involvement of ERK pathway in 3 gene mediated sex-specific phenotypes. The ERK pathway plays an important role in development, learning, and synaptic plasticity ^71, 72^. In addition, it has been shown that ERK signaling is involved in striatal behaviors ^37, 73-75^. We have observed reward-mediated hyperphosphorylated ERK1 in the dorsal striatum from 16p11.2 del/+ males, but not females consistent with the male-specific reward learning deficit in 16p11.2 del/+ mice ^10^, suggesting a role for ERK1 in striatum-dependent reward learning in males specifically. Therefore, we expect the connection between our 3 candidate genes and the ERK signaling pathway may provide an explanation for the molecular mechanisms of sex-specific striatal changes in 16p11.2 del/+ mice. This hypothesis is also supported by the strong connection between estrogen and the ERK signaling pathway. Estradiol influences memory consolidation via ERK signaling pathway in several brain regions ^76-79^. It has been shown that estrogen receptors interact with mGluR to activate ERK signaling ^80, 81^. Therefore, the connection between the 3 genes and ERK with estrogen may contribute to the sex-specific phenotypes of the 16p11.2 del/+ mice. Further studies will need to clarify the connection between ERK signal pathway and these 3 genes as an underlying mechanism of sex-specific phenotypes.

It has been reported that patients with 16p11.2 del syndrome show regional volumetric differences including accumbens, pallidum, caudate, and putamen, with increased FA in medial white matter ^12, 53, 82, 83^. In addition, 16p11.2 del/+ mice show increased volumes of several brain including striatum, nucleus accumbens, and globus pallidus ^48, 84, 85^. However, sex- and age-dependent brain structural changes in 16p11.2 deletion are not fully investigated. Previously, we have shown fiber tract changes in both male and female 16p11.2 del mice at 10 weeks of age ^31^. Here, we found decreased FA in 16p11.2 del/+ females compared to wt females at 6 weeks of age, however, not in males. These distinctive results indicate that FA changes in 16p11.2 del/+ mice are sex- and age-dependent, suggesting brain structural changes occur earlier in development in female 16p11.2 del/+ mice compared to male mice. In addition, the DTI results imply that the 3 gene hemi-deletion may not affect brain structural changes in the same way as 16p11.2 del. A previous study indicates that *Taok2* hemi-deletion mice show increased total brain volume compared with wt mice ^33^, however, 16p11.2 del/+ mice display decreased total brain volume ^48, 84^. These opposite brain structural changes between *Taok2* hemi-deletion mice and 16p11.2 del/+ mice may provide a hint to comprehend these distinctive brain structural changes between 3g del/+ mice and 16p11.2 del/+ mice. Uncovering the genetic mechanism of brain structural changes across development will require further study.

One important consideration for this study is whether the combinatorial effects of 3 genes on the behavioral phenotypes are the ‘sum of single gene effects’ or ‘combinatorial effects of 3 genes’. We believe it is the combinatorial effects of 3 genes based on previously described single gene mutation mouse studies. Although it has been shown that each gene (*Taok2, Sez6l2*, and *Mvp*) plays an important role in neuronal development as well as NDD ^33-35, 39-44, 86^, sex-specific phenotypes have not been observed in studies of single gene variant mouse lines. These studies suggest that the 3 gene hemi-deletion mediates male-specific phenotypes via combinatorial effects of 3 genes rather than just a summation of the effects of each gene. It is still possible that the combination of 2 genes may be sufficient to replicate certain phenotypes include male-specific phenotypes, so we cannot exclude the possibility that one out of the three genes is unnecessary to cause these effects. Here, we show that the 3 genes are sufficient to cause the male-specific phenotypes but not necessity of the 3 genes on the phenotypes. Therefore, further studies about necessity of each gene will be needed.

### Overall summary

This study demonstrates a novel approach to dissect large chromosomal regions to identify candidate sets of genes-of-interest and the characterization of a respective gene set in a novel experimental system carrying a selective hemi-deletion of three genes in the 16p11.2 region. 3g del/+ animals recapitulate male-specific striatal phenotypes observed in 16p11.2 del/+ mice. Changes in the expression levels of genes associated with ribosomal processing seem to constitute the transcriptomic basis for male-specific striatal vulnerability to NDDs. These findings could pave the way towards novel therapeutic and preventive strategies for NDDs.

## PUBLICATION ACKNOWLEDGEMENT

This work was supported by The University of Iowa Hawkeye Intellectual and Developmental Disabilities Research Center (HAWK-IDDRC) P50 HD103556 (T.A. and Lane Strathearn, PI), the Roy J. Carver Chair in Neuroscience (T.A.), Interdisciplinary Graduate Program in Genetics at University of Iowa (Y.V.), NIH grant R01 MH 087463 (T.A.), Simons Foundation Autism Research Initiative (SFARI) grant 345034 (T.A.), NIH grant T32 GM067795 (B.K.), Eagles Autism Challenge (T.N.-J.) and the Andrew H. Woods Professorship (T.N-J.).

Transgenic mice were generated at the University of Iowa Genome Editing Core Facility directed by William Paradee, PhD and supported in part by grants from the NIH and from the Roy J. and Lucille A. Carver College of Medicine. We wish to thank Norma Sinclair, Patricia Yarolem, Joanne Schwarting and Rongbin Guan for their technical expertise in generating transgenic mice.

The Neural Circuits and Behavior Core in the Iowa Neuroscience Institute provided equipment, facilities, and consultations services to support investigators in performing behavioral tasks. The Iowa Institute of Human Genetics provided Sanger DNA sequencing and RNA sequencing services. The Iowa Magnetic Resonance Research Facility provides access to a small animal MRI scanner as well as the necessary data processing equipment for animal MR imaging.

