## Supplemental figures for "Dissecting 16p11.2 hemi-deletion to study sex-specific striatal phenotypes of neurodevelopmental disorders"

**Supplementary Figure 1. Gene modification design**

**A** Wildtype sequence (**TAO kinase 2** **Exon 2**):

TACCTACCTTTCCTCAGCAGTACCTCCCCCTCATCAGTAAATGAGGGGAGGAGGAGGTAGAAACCTGAGGGA
GGACCTCTCTCTCCCTCTGGGCCCTATCTTAGCTCTAAGGGTCCTATGTCCTTTTTCCAG**GCAAGATCCCAATC**
**TCAGGGCCCCCTGGGGCCATCATGCCAGCTGGGGGCCGGGCCGGGAGCCTGAAGGACCCTGATGTGGCT**
**GAGCTCTTCTTCAAGGATGACCCTGAGAAGCTCTTCTCTGACCTCCGGGAGATCGGCCATGGCAGCTTTGG**
**AGCAGTGTACTTT**GTGAGTTGGGTCTTGAAAAGGGTAAAGCAGGGCTCAGTCTCTTTCAACCTGTGGGTCT
CCAGGCCTCTGCACCACTCCACCAAAATAATCCTTCCCACCCTCTTCTAATAGCTCAGCGGGTCCTCTTTCACC
CCATGCCCAAGGTGGTCTTTTCCATCCTCCAATCTGGTCCTCTAGGCCCGGGATGTCCGGAACAGTGAGGTG
GTGGCCATCAAGAAGATGTCCTATAGTGGGAAGCAATCAAATGAGGTGAGTCAGGTTGATTAACATCAGGTT
GTGGAGGG

Mutant Sequence:

TACCTACCTTTCCTCAGCAGTACCTCCCCCTCATCAGTAAATGAGGGGAGGAGGAGGTAGAAACCTGAGGGA
GGACCTCTCTCTCCCTCTGGGCCCTATCTTAGCTCTAAGGGTCCTATGTCCTTTTTCCAG**GCAAGATCCCAATC**
**TCAGGGCCCCCTGGGGCCATCATGCCAGCTGGGGGCCGGGCCGGGAGCTCAGCCACTxxxxxxxxxxCTGA**
**GCTCTTCTTCAAGGATGACCCTGAGAAGCTCTTCTCTGACCTCCGGGAGATCGGCCATGGCAGCTTTGGAG**
**CAGTGTACTTT**GTGAGTTGGGTCTTGAAAAGGGTAAAGCAGGGCTCAGTCTCTTTCAACCTGTGGGTCTCCA
GGCCTCTGCACCACTCCACCAAAATAATCCTTCCCACCCTCTTCTAATAGCTCAGCGGGTCCTCTTTCACCCCAT
GCCCAAGGTGGTCTTTTCCATCCTCCAATCTGGTCCTCTAGGCCCGGGATGTCCGGAACAGTGAGGTGGTGG
CCATCAAGAAGATGTCCTATAGTGGGAAGCAATCAAATGAGGTGAGTCAGGTTGATTAACATCAGGTTGTGG
AGGG

**TCAGCCACT – inversion of part of the deleted sequence**
**xxxxxxxxxx – 10 bp deletion = out of frame mutation; premature stop in exon 4**

**B** Wildtype sequence (**seizure related 6 homolog like 2 (Sez6l2)** **Exon 2** **ATG in Exon 1**):

AGGATGTAGAGGAATGGAGAGGTTCACAGGACTTCACCTCCTAG**GTCTGCCCCTGAAGGAGGATGAGATG**
**ATGCCAGAGCCTGGAAGTGAGACTCCCACAGTGGCCTCTGAGGACCTGGCTGAGCTGCTCCATGGGGCTT**
**TGCTGCGGAAGGGCCCAGAGATCGGCTTCTTGCCGG**GTGAGGCCCACAGTGTGGCATAGGAGTAGAGAG
AGAGGCTATGTCGCTGAGAGCTGGAGTGCCTGGCTAGAGGGAAGGCGGGTTAGAGAGAGTTCTGTGGGA
GAGACCCCCTAGGAAGCTGAGAAAGAGTCAAAGCTGGCC

Mutant Sequence:

AGGATGTAGAGGAATGGAGAGGTTCACAGGACTTCACCTCCTAG**GTCTGCCCCTGAAGGAGGATGAGATG**
**ATGCCAGAGCCTGGAAGTGAGACTCCCACAGTGGCCTCTGAGGACCTGGCTGAGCTGCTCCATGGGG::::::**
**:::::::::::CCAGAGATCGGCTTCTTGCCGG**GTGAGGCCCACAGTGTGGCATAGGAGTAGAGAGAGAGGCTATG
TCGCTGAGAGCTGGAGTGCCTGGCTAGAGGGAAGGCGGGTTAGAGAGAGTTCTGTGGGAGAGACCCCCTA
GGAAGCTGAGAAAGAGTCAAAGCTGGCC
**::::::::::::::::: – 17 bp deletion = out of frame mutation; premature stop in exon 3**

C

Wildtype sequence (**major vault protein (Mvp)** Exon 2; *Exon 3;* **ATG in Exon 1**):

**TCACCATGGCAACTGAAGAGGCCATCATCCGCATCCCCCCATACCACTACATCCATGTGCTGGACCAGAACA GTAATGTGTCCCGTGTAGAGGTTGGACCAAAGACCTACATCCGGCAGGACAATGAGAG**GTTGGTGTAGAG CTGTCCCAGCCTGGCTGGTGGGAATGACCCTCATCTGGGTGGCCGGGAGGTTTCTCTTGCTTTTACTGTCTCC TTTGGAACATCATCCTGGCTCCTCACGCCCCTTCTTATCTTACAACAG***GGTACTGTTTGCCCCAGTTCGCATGG TGACGGTCCCACCACGCCACTACTGCATAGTGGCCAACCCTGTGTCCCGGGACGCCCAGAGTTCTGTGTTGT TTGACGTCAC***

Mutant Sequence:

**TCACCATGGCAACTGAAGAGGCCATCATCCGCATCCCCCCATACCACTACATCCATGTGCTGGACCAGAACA GTAATGTGTCCCGTGTAGAGGTTGGACCAAAGACCTACTATCCGGCAGGACAATGAGAG**GTTGGTGTAGA GCTGTCCCAGCCTGGCTGGTGGGAATGACCCTCATCTGGGTGGCCGGGAGGTTTCTCTTGCTTTTACTGTCT CCTTTGGAACATCATCCTGGCTCCTCACGCCCCTTCTTATCTTACAACAG***GGTACTGTTTGCCCCAGTTCGCAT GGTGACGGTCCCACCACGCCACTACTGCATAGTGGCCAACCCTGTGTCCCGGGACGCCCAGAGTTCTGTGTT GTTTGACGTCAC***

**T – 1 bp insertion = out of frame mutation; premature stop (TGA) in exon 2**

**Supplementary Figure 2. Expression of genes adjacent to modification in 3g del/+ mice**

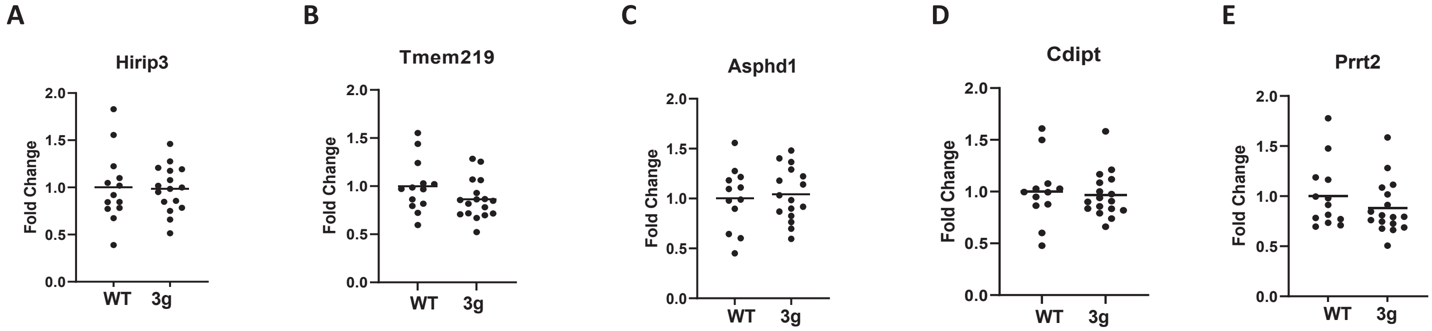

**Supplementary Figure 3. Demographics of 3g del/+ mice inheritance and growth curve**

**A**

#### 3g del/+ male mouse x wt female mouse

| | WT (number of animals) | 3g del/+ (number of animals) | 3g del/+ inheritance (%) |
|---|---|---|---|
| **Male** | **62** | **58** | **48.33** |
| **Female** | **47** | **44** | **48.35** |

**B**

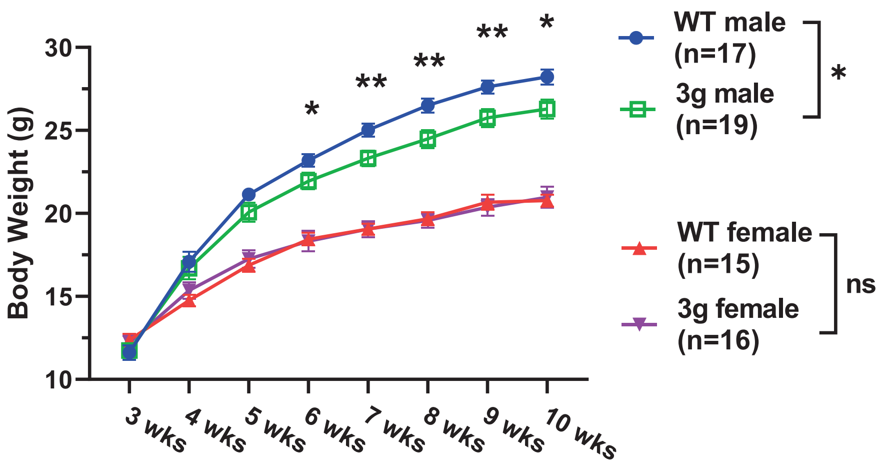

Supplementary Figure 4. Rotarod test on 3g del/+ mice

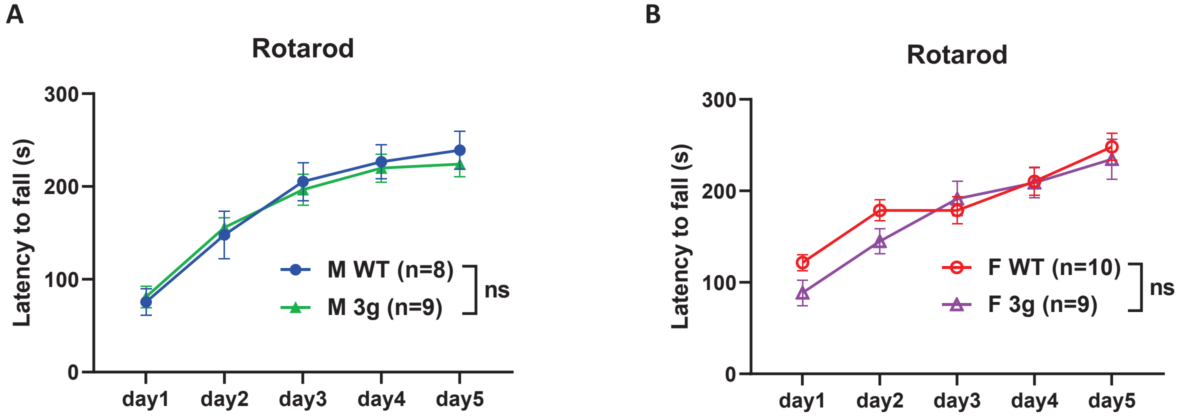

#### Supplementary Figure 5. Body weight during food restriction

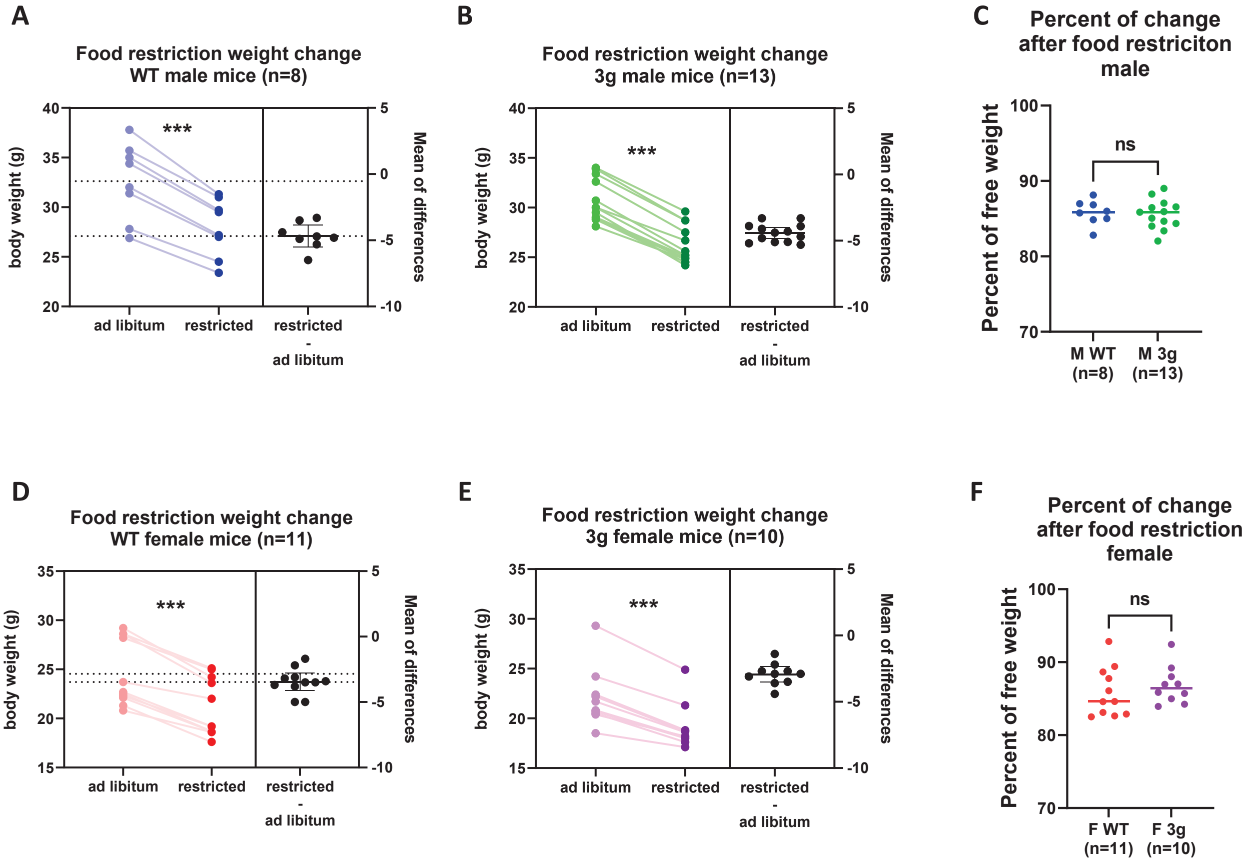

### Supplementary Figure 6. Additional behavioral phenotypes of 3g del/+ mice

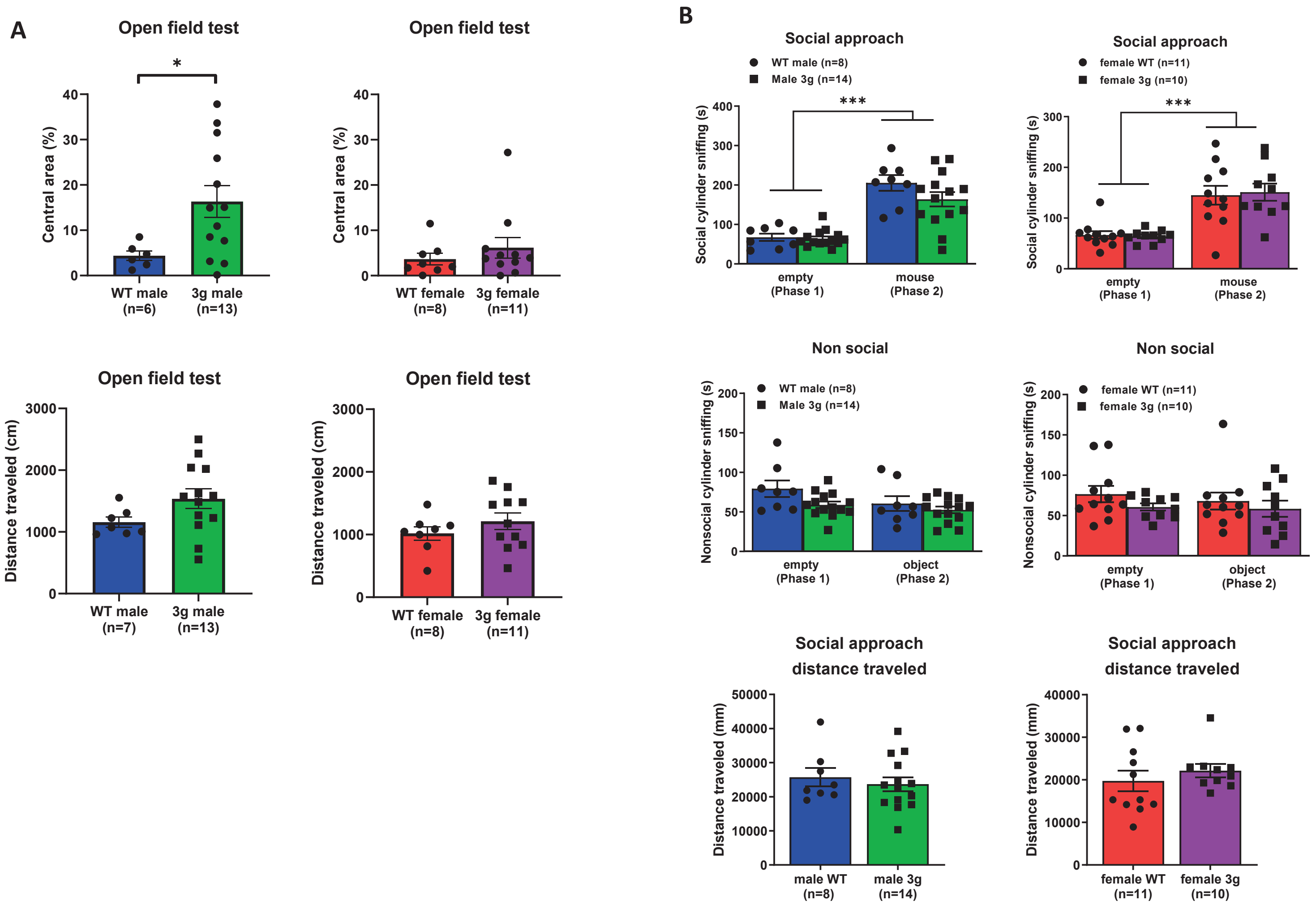

**Supplementary Figure 7. Quadrant plot of 248 DEGs with labelled ribosomal genes**

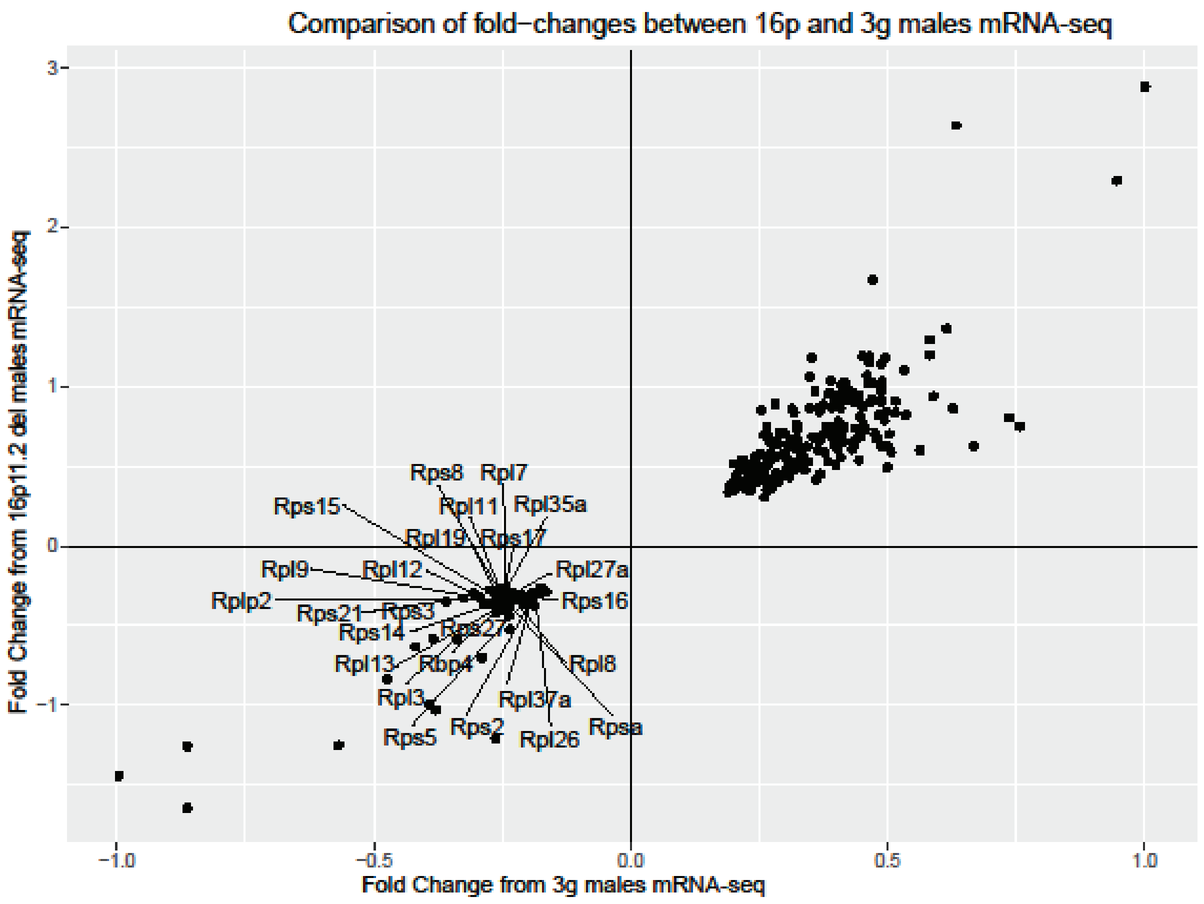

**Supplementary Figure 8. Pathway analysis of RNAseq results on 3g males (569 DEGs)**

A

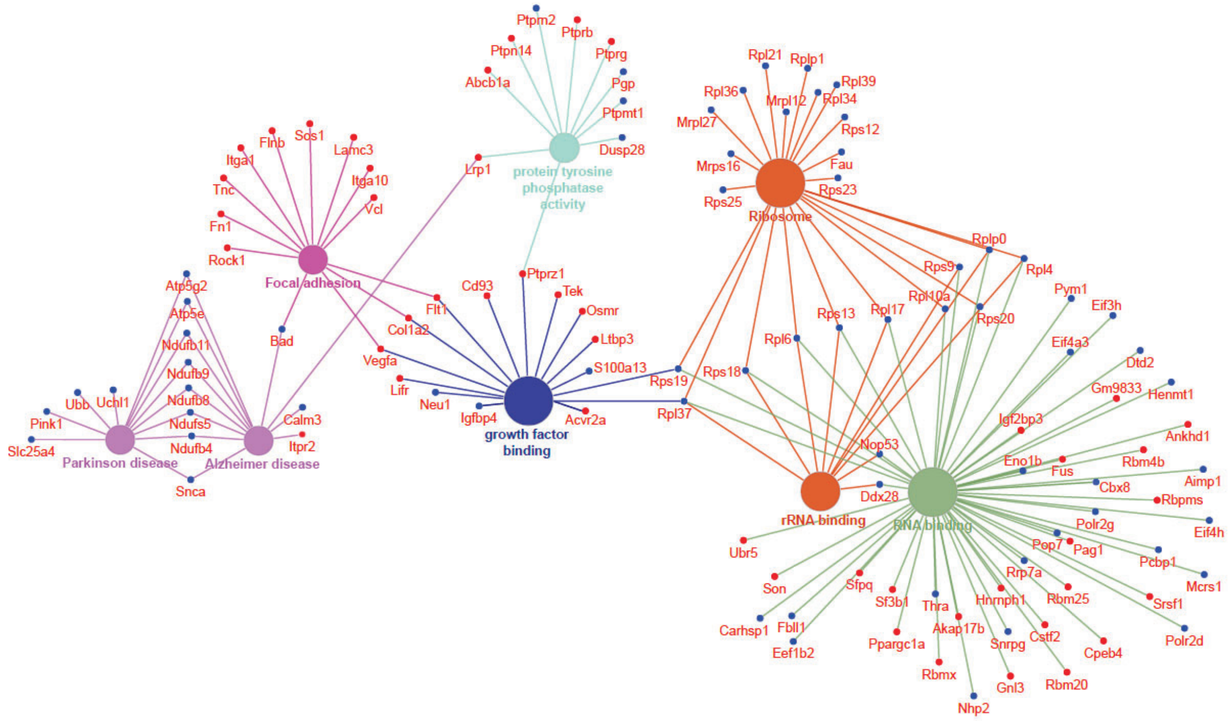


B

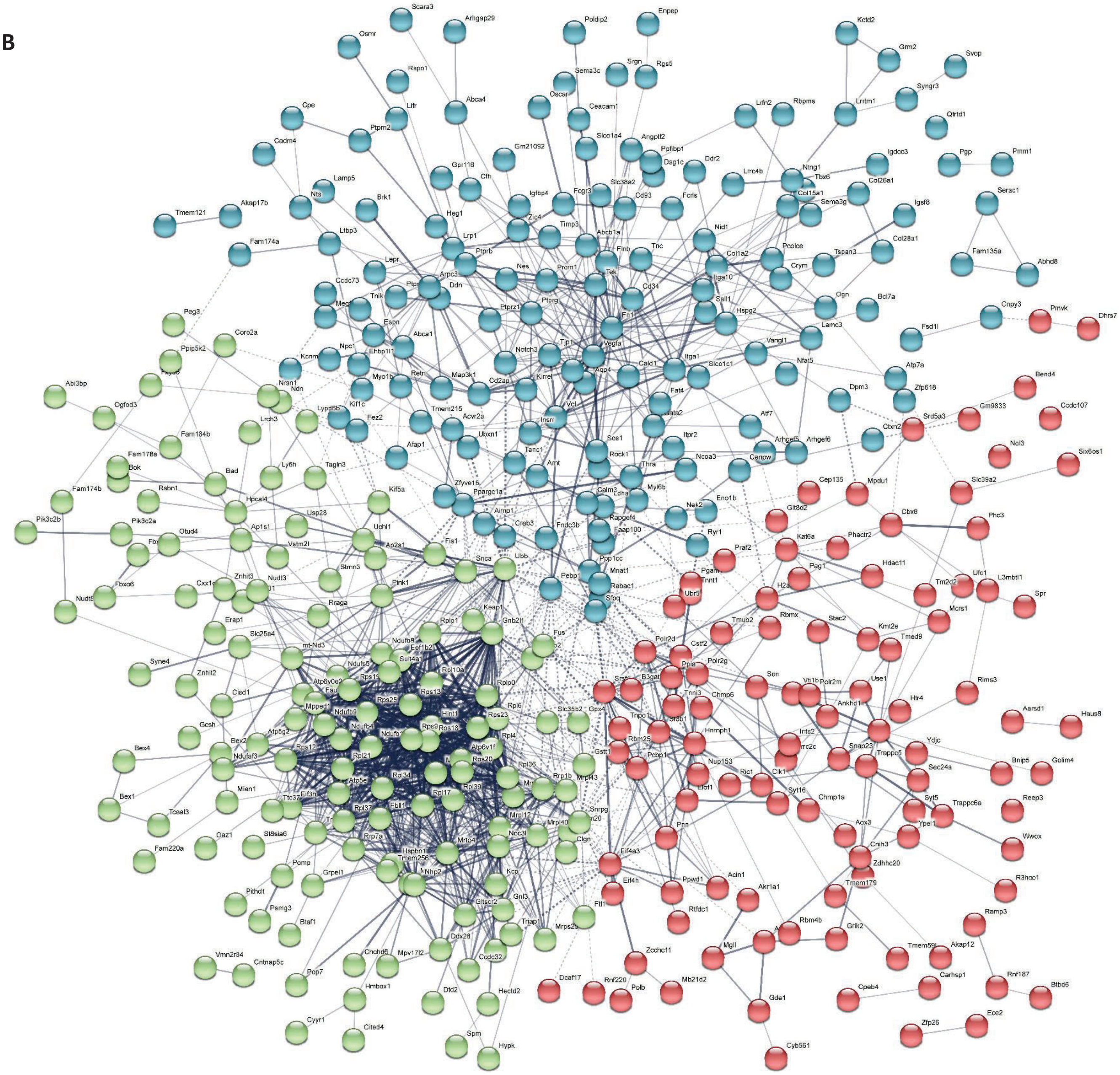

**Supplementary Figure 8. Pathway analysis of RNAseq results on 3g males (569 DEGs)**

C

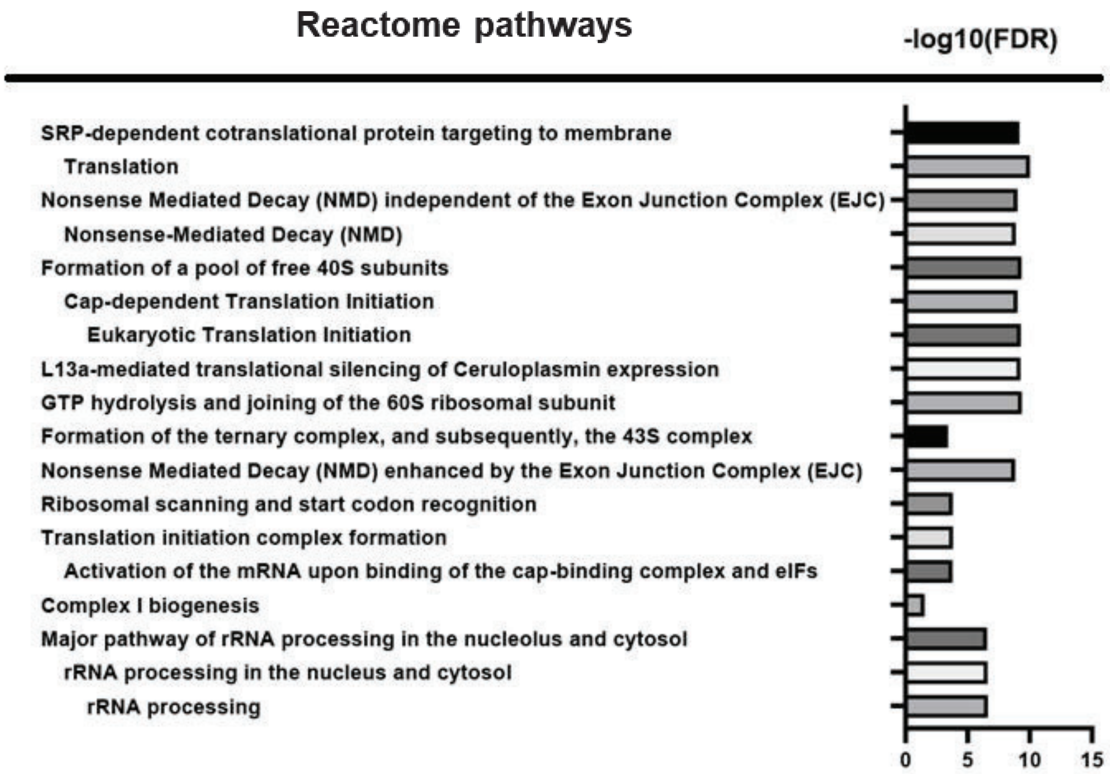

**Supplementary Figure 9. Pathway analysis of RNAseq results on 16p11.2 males (633 DEGs)**

A

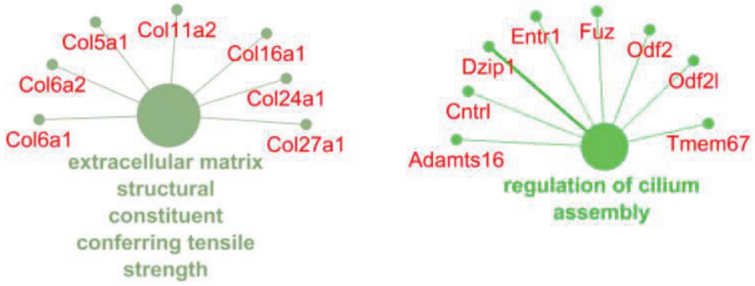


B

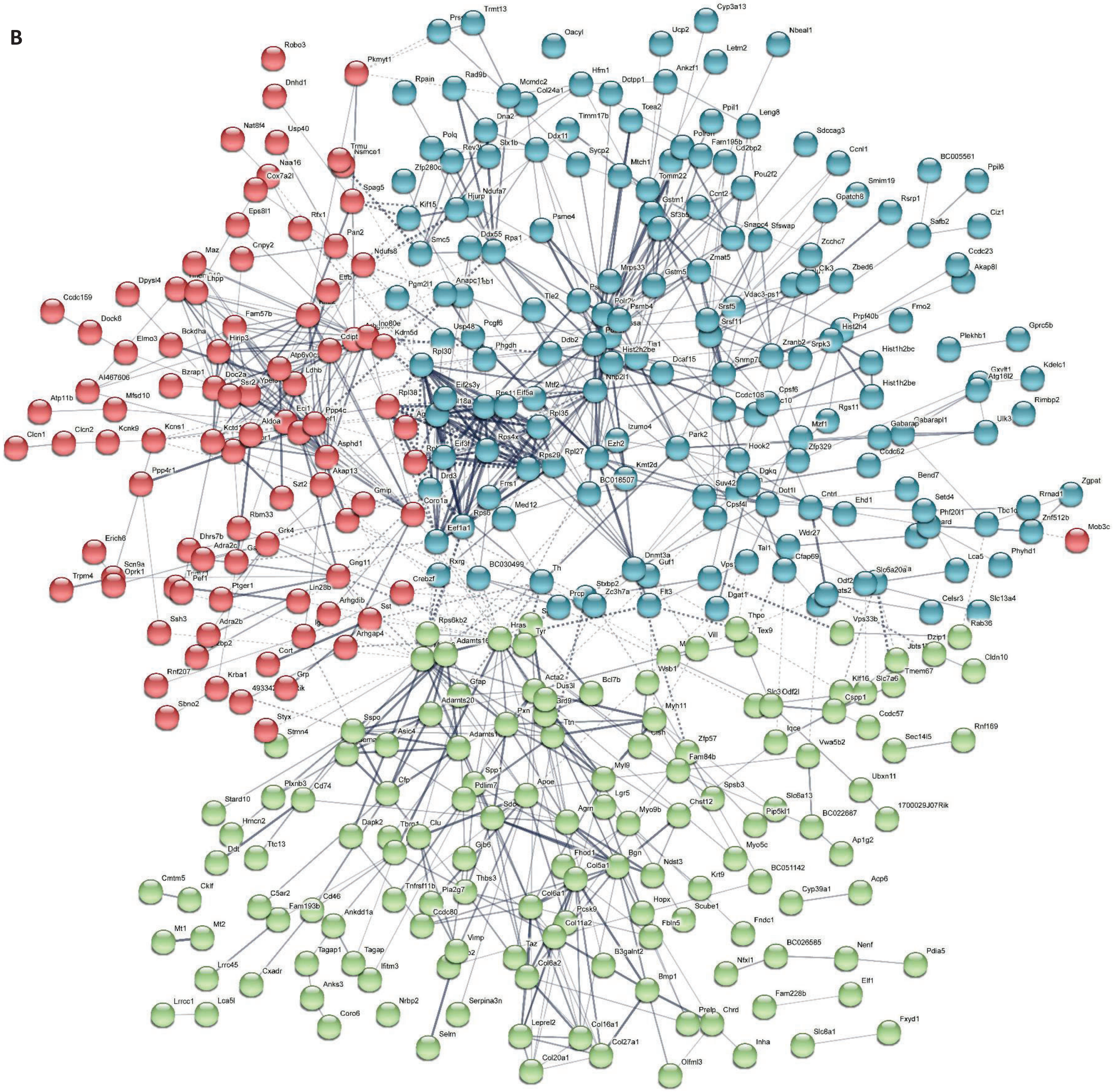

**Supplementary Figure 9. Pathway analysis of RNAseq results on 16p11.2 males (633 DEGs)**

C

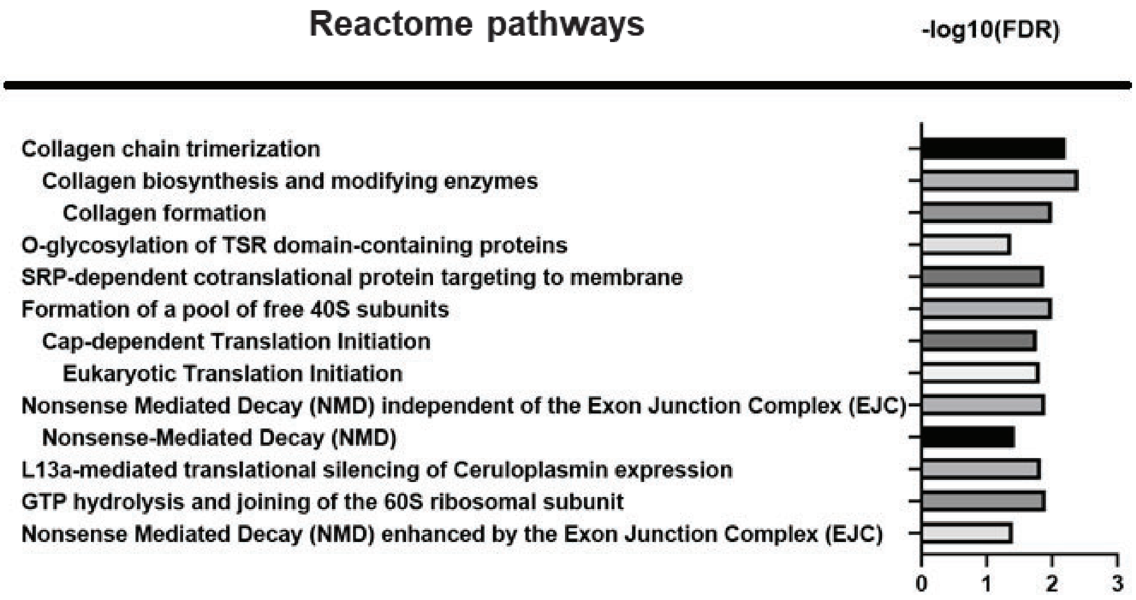

**Supplementary Figure 10. Pathway analysis of RNAseq results on 3g females (340 DEGs)**

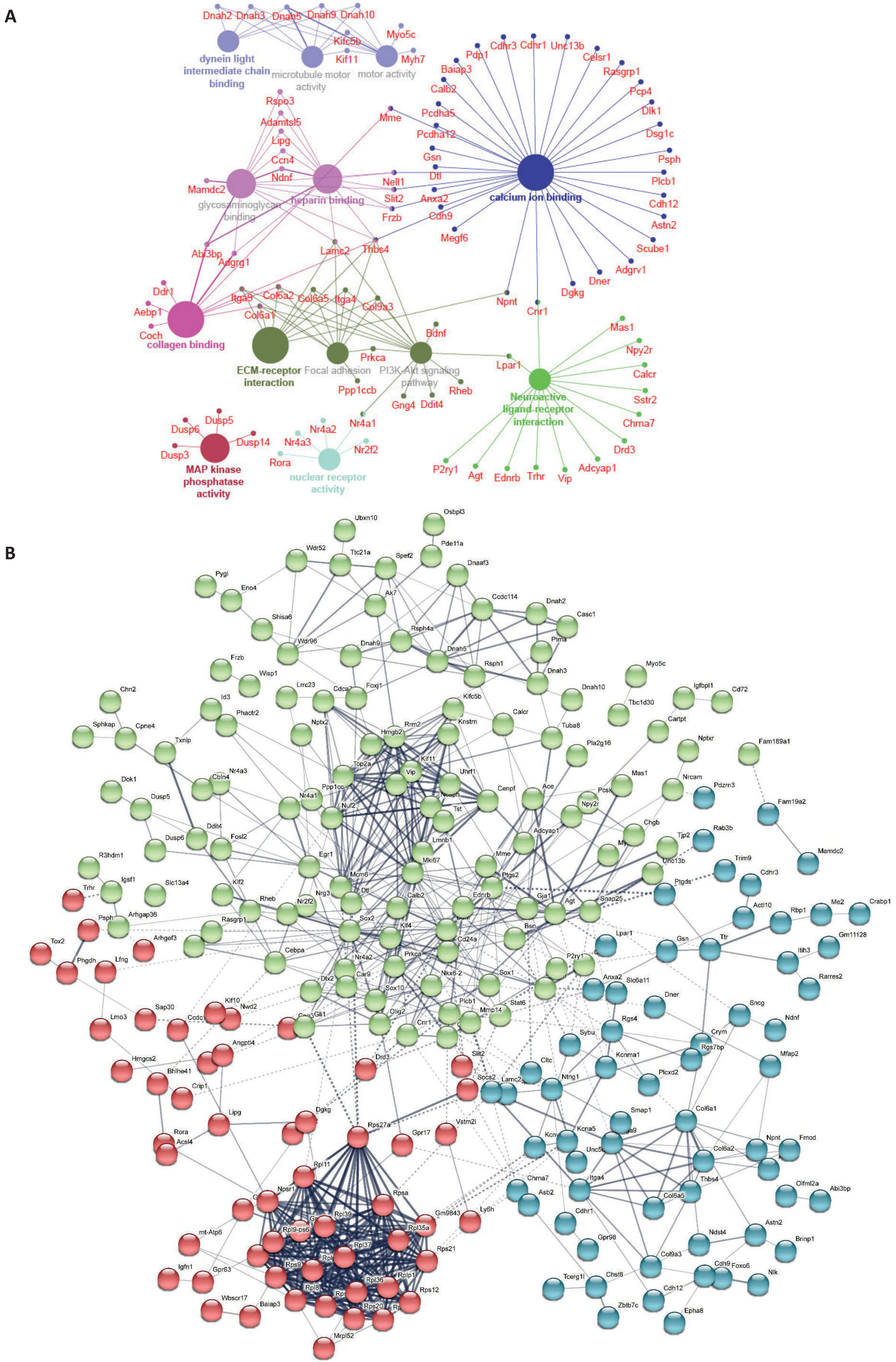

**Supplementary Figure 10. Pathway analysis of RNAseq results on 3g females (340 DEGs)**

C

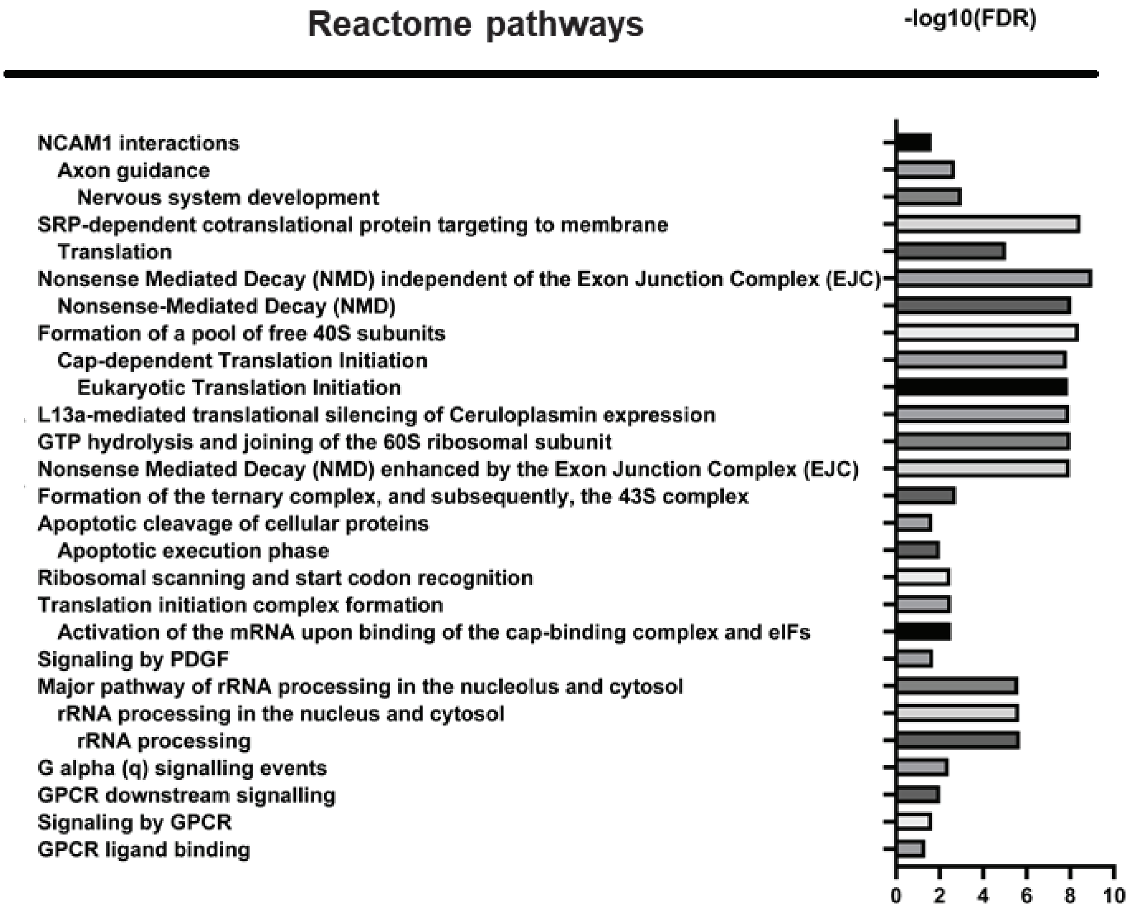

**Supplementary Figure 11. Pathway analysis of RNAseq results on 16p11.2 females (121 DEGs)**

A

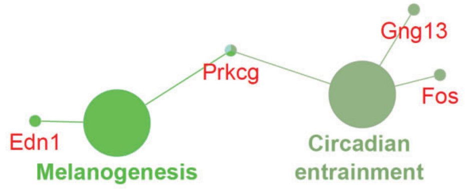


B

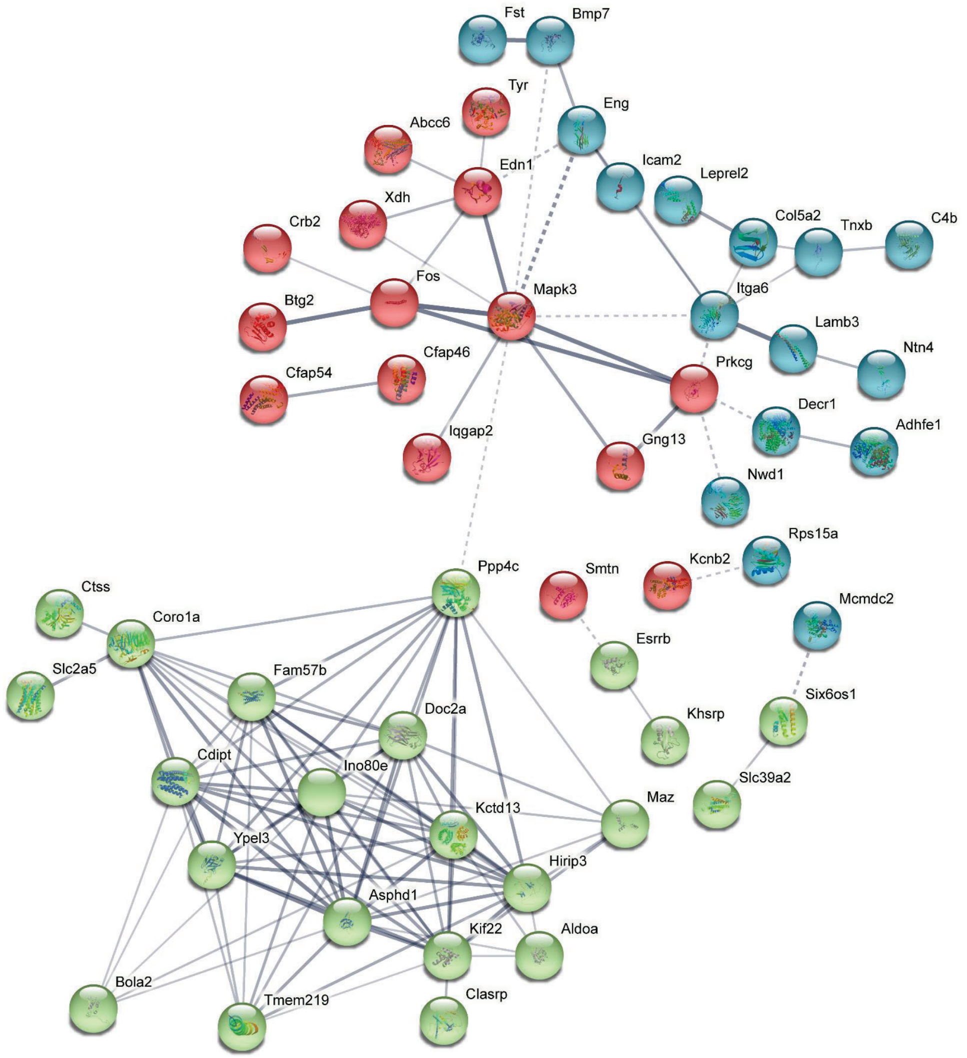


C

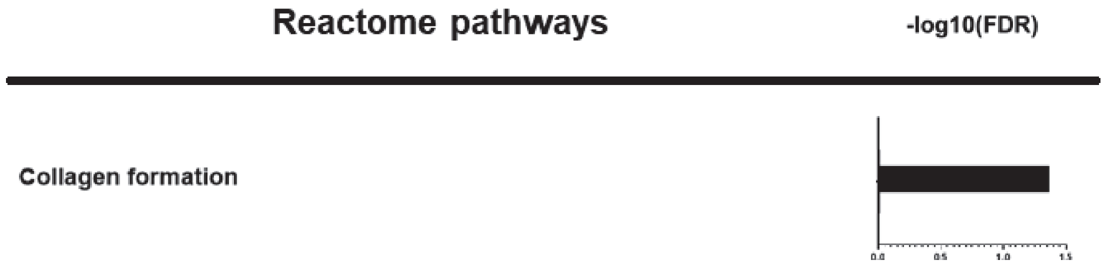

**Supplementary Figure 12. Fiber tract changes in 3g mice compared to wild types**

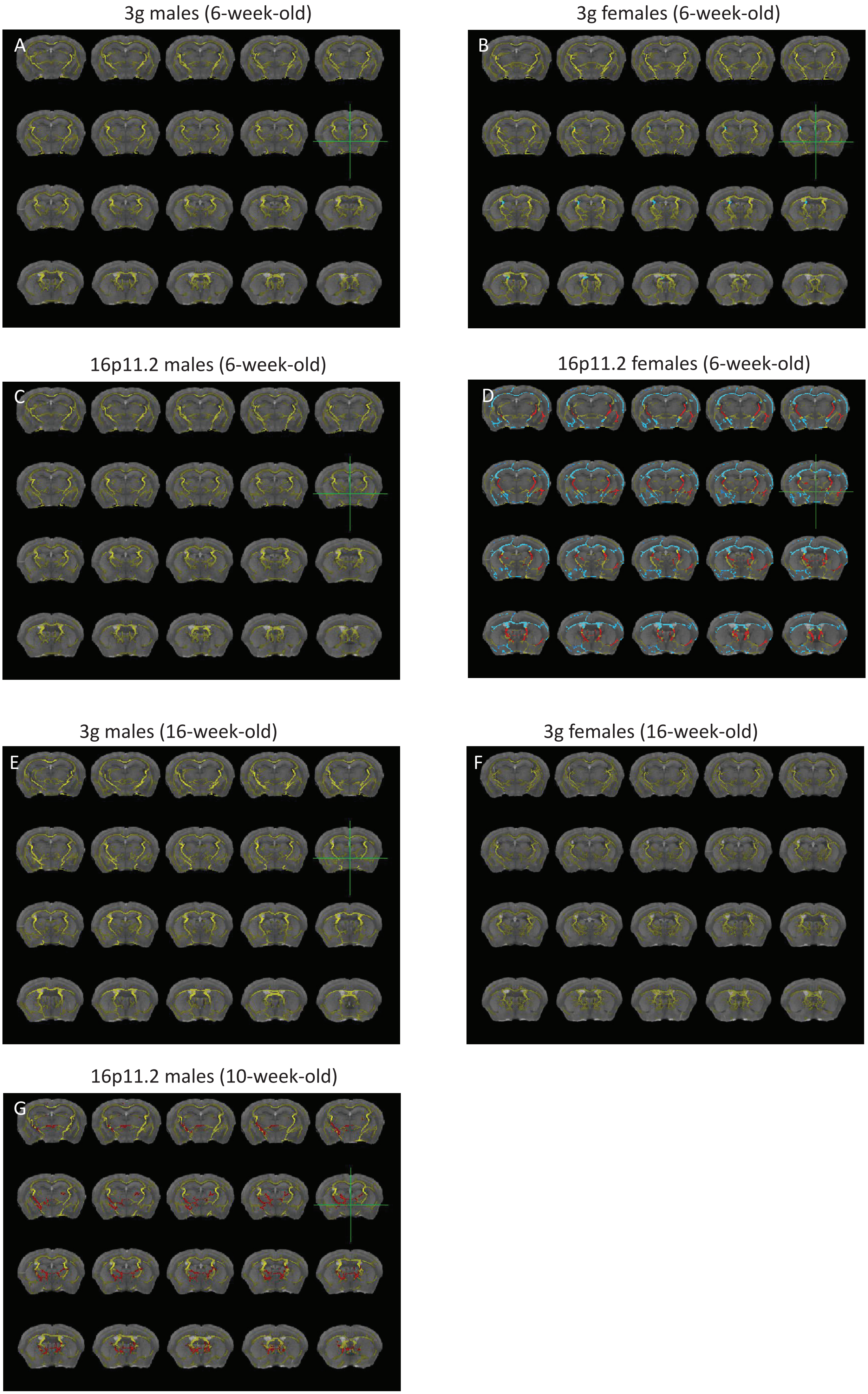

**Supplementary Figure 13. Graphic abstract**

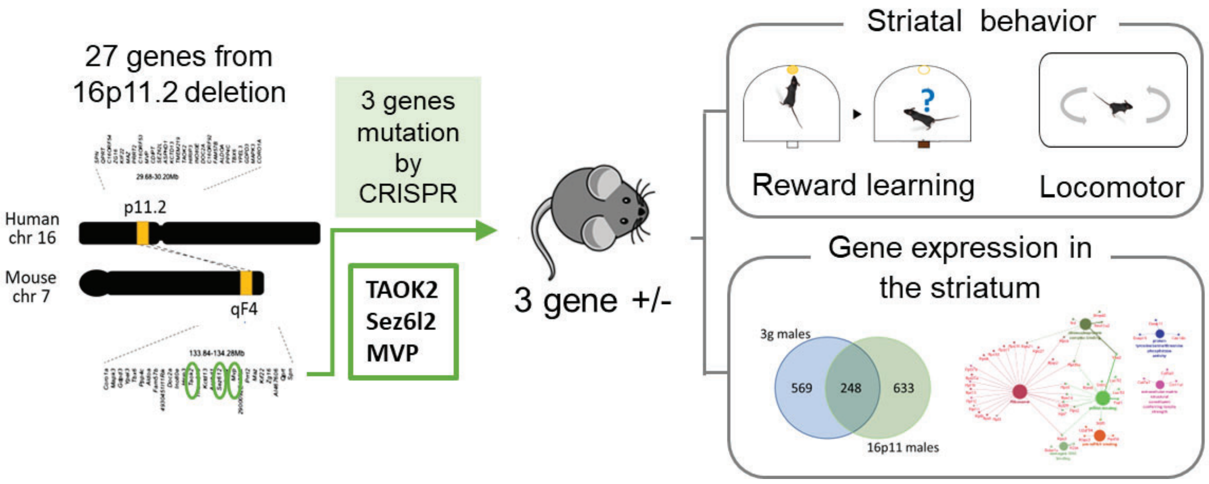
