## Supplemental method for "Dissecting 16p11.2 hemi-deletion to study sex-specific striatal phenotypes of neurodevelopmental disorders"

METHODS AND MATERIALS

Animals

All procedures were approved by the University of Iowa Institutional Animal Care and Use Committees (IACUC), and followed policies set forth by the National Institutes of Health Guide for the care and use of laboratory animals. Mice were housed in groups (2-5 per cage) in a room with a 12-hour light/dark cycle, and were allowed ad libitum access to food and water. Male and female mice were used in all experiments. 3g del/+ mice were generated at Genome Editing Core of Carver College of Medicine at University of Iowa by CRISPR/Cas9-induced gene modifications and backcrossed to C57BL6/J. Male 3g del/+ mice were bred with S129 from The Jackson Lab (#002448) to generate experimental cohorts of littermates, to match mixed genetic background of 16p11.2 del/+ mice. 16p11.2 del/+ mice were purchased from The Jackson Lab (#013128) and breed with B6129SF1/J females (#101043), as previously studies [1-3]. Experimental data were collected from littermate animals during the same time period. Mice aged between 10 and 16 weeks were used for experiments unless otherwise specified.

Generation and genotyping of 3g del/+ mice

*Preparation of* *Cas9 RNPs and the electroporation mix*

C57BL/6J mice were purchased from Jackson Labs (000664; Bar Harbor, ME). Male mice older than 8 weeks were used to breed with 3-5 week-old super-ovulated females to produce zygotes for electroporation. Female ICR (Envigo; Hsc:ICR(CD-1)) mice were used as recipients for embryo transfer.

Chemically modified CRISPR-Cas9 crRNAs and CRISPR-Cas9 tracrRNAs were purchased from IDT (Alt-R® CRISPR-Cas9 crRNA; Alt-R® CRISPR-Cas9 tracrRNA (Cat# 1072532)). The crRNAs and tracrRNA were suspended in T10E0.1 and combined to 1 ug/ul (~29.5 uM) final concentration in a 1:2 (ug:ug) ratio. The RNAs were heated at 98C for 2 min and allowed to cool slowly to 20C in a thermal cycler. The annealed cr:tracrRNAs were aliquoted to single-use tubes and stored at -80C.

Cas9 nuclease was also purchased from IDT (Alt-R® S.p. HiFi Cas9 Nuclease). Individual Cr:tracr:Cas9 ribonucleoprotein complexes were made by combining Cas9 protein and cr:tracrRNA in Opti-MEM (Gibco; 31985062; final concentrations: 916 ng/ul (~5.7 uM) Cas9 protein and 366 ng/ul (~11.2 uM) cr:tracrRNA). The Cas9 protein and annealed RNAs were incubated at 37C for 15 minutes. The three RNP complexes were mixed resulting in final concentrations of 916 ng/ul (~5.7 uM) Cas9 protein and 122 ng/ul (~3.7 uM) each cr:tracrRNA.

Tao_Guide_A	AGAAGAGCTCAGCCACATCA

Sez_Guide_B	CATGGGGCTTTGCTGCGGAA

Mvp_Guide_A	TTGGACCAAAGACCTACATC

*Collection of embryos and electroporation*

Pronuclear-stage embryos were collected using methods described in [4]. Embryos were collected in KSOM media (Millipore; MR101D) and washed 3 times to remove cumulous cells. The embryos were treated for 10 seconds in acid tyrodes solution (Sigma; T1788) to weaken the zona. The embryos were then washed 5 times with Opti-MEM and 25-30 were aligned between the electrodes of a slide glass plate electrode with a 1 mm gap and platinum leads. The acid-treated embryos were overlayed with the electroporation mix and electroporated for 7 pulses at 30V (3 msec on:100 msec off). Electroporation was performed using an ECM830 square wave electroporator (BTX). Embryos were immediately implanted into pseudo-pregnant ICR females.

*Genotyping*

PCR-genotyping and Sanger sequencing was done to identify founders with the intended knock-in mutation.

Tao_101	TACCTACCTTTCCTCAGCAGTA

Tao_102	CCCTCCACAACCTGATGTTAAT

WT:	583 bp 🡪 Hpy188I 🡪 330 + 253

Mutation:	573 bp 🡪 Hpy188I 🡪 330 + 39 + 204

Sez_101	AGGATGTAGAGGAATGGAGAGG

Sez_102	GGCCAGCTTTGACTCTTTCT

WT:	317 bp 🡪 Apa1 🡪 155 + 162

Mutation:	300 bp 🡪 Apa1 🡪 300

Mvp_101	TCACCATGGCAACTGAAGAG

Mvp_102	GTGACGTCAAACAACACAGAAC

WT:	371 bp

Mutation:	372 bp

Behavioral tasks

*Activity monitoring*

Mice locomotor activity was measured by activity monitoring using an infrared beam-break system (Opto M3, Columbus Instruments, Columbus, OH) as previously described [1, 5]. To minimize the potential impact of novel environment, mice were habituated in the activity chambers for 1 week before the data collection. The data was achieved for 1 week after the habituation week. ANOVAs were used with genotype (wt or 3g mice) as the between-subjects factor and time as the within-subjects factor, as previously described [1, 5], using GraphPad 9 (La Jolla, CA).

*Rotarod task*

Motor coordination ability and motor learning were evaluated using a rotarod. Mice were placed on a rotating rod, and the time at which each mouse falls off or makes one complete backward rotation was documented. The rotarod was set at accelerating speed (4 to 40 rpm). Each day, mice were tested 3 trials for up to 5 minutes trial for 5 days (total 15 trials). The latencies of each trial were averaged over the 1 day of data collection. Inter-trial intervals were 10-20 min.

*Operant task*

Mice were allowed to acclimate to a 0900–2100 hours reversed light cycle one week prior to obtain food-restricted weights of 85-90% of free-feeding weights. Mice had unlimited access to water. To minimize the chance of food related competition, the mice were housed in cages of one to two littermates per cage. If mice dropped below 85% of their free-feeding weight or signs of aggression were noted between cagemates, mice were single housed for the remainder of the experiment. Chocolate flavored Ensure Original Nutrition Shake (Abbott) diluted to 50% with water was used as a reinforcer. Animals were exposed to the reinforcer in small dishes for the three days prior to the start of the experiment to prevent a neophobic response to the reinforcer in the experiment. During this time, the consumption of each mouse was observed.  Mice were trained in Mouse 9 Hole Operant Test Chambers (Lafayette Instruments, Lafayette IN USA) with the holes/operanda (containing lights and infrared beams to register responses) in the rear and the liquid reward magazine (also containing a light and IR beam) in the front. Only the middle nose-poke hole was accessible to reduce extraneous responding to non-active holes. The first day of operant training consisted of magazine training: one 20uL reward was delivered (signaled by the illumination of the reward receptacle), once the reward was collected, a variable inter-trial interval (vITI) (average 5s, range: 3, 4, 5, 6, 7) was initiated after which the next reward was delivered. The session ended after 30 rewards were delivered or 30 minutes elapsed. Next, mice underwent two days of autoshaping: after an ITI (vITI 60, average 60s, range: 30, 45, 60, 75, 90), the light in the center nose-poke hole was illuminated for 8 seconds, if the mouse did not respond, a reward was delivered after the hole light turned off. If the mouse responded while the hole light was on, a reward was immediately delivered. After the reward was collected, another ITI was initiated before the next trial commenced. A session ended after 30 minutes. Mice were then moved onto a fixed ratio (FR) schedule of reinforcement where, after an ITI (vITI 60), one nose-poke response to the lit center hole was required to earn a reward (FR1), as previously described [2, 6]. FR sessions ended after 30 minutes. FR1 training continued for eight days. Responses per day for each animal were recorded and analyzed using a repeated measures ANOVA with genotype as the between-subject factors and day as the within-subjects factor, as previously described [2], using GraphPad 9. One week after FR1, the mice were transitioned onto a progressive ratio (PR) schedule of reinforcement as previously described [2, 6]. In PR, the response requirement increased in an arithmetic sequence (1, 2, 4, 7, 11, 16,…n) every third trial. PR testing was terminated after no nose-poke was made for five minutes or after one hour. The PR breakpoint was defined as the last trial where the response requirement was completed and reinforcement was delivered. Average breakpoints were assessed by unpaired t-tests using GraphPad 9.

*Open field task*

Open field activity was recorded by EthoVision video tracking system (Noldus Information Technology), measuring peripheral, center, and total distance, and time spent in the center. As previously described [3], mice were familiarized with the procedure room with a 30-min acclimation. then individual mice were placed in a clear 35x35x45 cm Plexiglas arena (SanDiego Instruments) for 10 min trial. In-between trials, the area was cleaned with 70% ethanol.

*Social approach task*

Social approach task was recorded by EthoVision video tracking system (Noldus Information Technology), using an IR-compatible digital camera, measuring the distance traveled and time of cylinder exploration. As previously described [3], the testing area was a 3-chambered arena, a bottomless black Plexiglass rectangular box (16 long x 6 wide x 9 inches tall), with two end chambers of equal size (5.9 x 6 inches) and a smaller middle chamber (4.2 x 6 inches). The procedure was performed in a dark room (2-5 lux) with an infrared light panel which is located below the arena. At the start of task, mice were habituated to the arena with two identical clear Plexiglas cylinders (each 3-inch diameter, 5.8-inch tall) by allowing them to explore for 10 min (phase 1). Then, social stimulus mouse, which is gonadectomized A/J mouse (Jackson Laboratories), was placed in one cylinder and a novel object was loaded in the other cylinder. The test mouse was allowed 10 min to freely explore the arena and cylinders containing the object or mouse (phase 2).

RNA quantification

RNA isolation was conducted as previously described [2]. Mice were cervically dislocated and decapitated, and the total striatum was dissected bilaterally using a mouse brain matrix allowing 1 mm sections. The tissue was stored in RNAlater, then RNA was extracted using Qiagen RNeasy kit (Qiagen) according to the manufacturer's instructions. cDNA was synthesized using the iSript cDNA Synthesis Kit (Bio-Rad). Quantitative real time polymerase chain reaction (qPCR) was performed in The Applied Biosystems® QuantStudio™ 7 Flex Real-Time PCR System (Thermo Fisher Scientific) using the Fast SYBR Green RT-PCR kit (Thermo Fisher Scientific) according to the manufacturer's instructions. The specific RNA expression was calculated using GraphPad Prism 9 (San Diego, CA, USA) based on the comparative Ct method, normalized to the geometric mean of GAPDH and Tubulin.

Primer information

| *Asphd1* | Forward | TCCCTCCTGGTTGTGAGTTG |
| --- | --- | --- |
|  | Reward | GTGTGCAGGAAGGAGTCGTC |
| *Cdipt* | Forward | TTGTCTTCGCCATCATTTCC |
|  | Reward | TCCATCGAAAGCGTCTAGGA |
| *Gapdh* | Forward | TGCACCACCAACTGCTTA |
|  | Reward | GGCATGGACTGTGGTCATGA |
| *Hirip3* | Forward | AGGGAAGAGAGTGGGAGCAG |
|  | Reward | TCTCCTCCTGGCAGTTGACA |
| *Mapk3* | Forward | CCAAAGCTCTTGACCTGCTG |
|  | Reward | CCAGCGCTTCCTCTACTGTG |
| *Mvp* | Forward | ATGTCAAGACGGGAAAGGTG |
|  | Reward | TTTCCCACAGGACTTCATCC |
| *Prrt2* | Forward | ACACCCAGTCAGACCCTCAG |
|  | Reward | TCAGGACCTCTGTGGTAGGG |
| *Sez6l2* | Forward | TGTGAGCGTGACAGACTTGC |
|  | Reward | GGGTCTCCTCACCCTGTAGC |
| *Taok2* | Forward | AGGAAGTGCGGTTCTTACAGA |
|  | Reward | GAGCCCAGGCAATACTCCAT |
| *Tmem219* | Forward | CCACTTTGATCCTGCTGCTC |
|  | Reward | GTTGTGTGGGTGAGGAGGAA |
| *Tubulin* | Forward | ATGCGCGAGTGCATTTCA |
|  | Reward | CACCAATGGTCTTATCGCTGG |

RNA-seq

*Library preparation*

RNA quality was assessed using a BioAnalyzer (Agilent) and samples with RNA integrity number (RIN) >8 were used to perform RNA-seq. RNA library preparation from WT mice (n = 8 samples, 4 males and 4 females) and 3g del/+ mice (n = 8 samples, 4 males and 4 females) for 3g del/+ mice and WT mice (n = 8 samples, 4 males and 4 females) and 16p11.2 del/+ mice (n = 8 samples, 4 males and 4 females) for 16p11.2 del/+ mice were prepared at the Iowa Institute of Human Genetics (IIHG), Genomics Division, using the Illumina TruSeq Stranded Total RNA with Ribo-Zero gold sample preparation kit (Illumina, Inc., San Diego, CA). Library concentrations were measured with KAPA Illumina Library Quantification Kit (KAPA Biosystems, Wilmington, MA). Pooled libraries were sequenced on Illumina NovaSeq 6000 sequencers with 100-bp Paired-End chemistry (Illumina) at the IIHG core, resulting in an average of 98,575,000 reads per sample in WT mice and 99,175,000 reads per sample in het mice. The dataset supporting the conclusions of this article is available in the NCBI’s Gene Expression Omnibus repository, GEO Series accession GSE224750.

*RNA-seq analysis*

Sequencing data was processed with the bcbio-nextgen pipeline (https://github.com/bcbio/bcbio-nextgen). In brief, raw reads were aligned to the Mus musculus reference genome mm10 (mus_musculus_vep_100_GRCm38) with STAR (version 2.6.1d)[7]. MultiQC was then used for quality control and assurance analysis of the resulting bam file by comparison to metrics gathered from bcbio-nextgen, samtools (version 1.9), and fastqc (version 0.11.8) [8, 9]. Quantified reads were assigned to genes (features) annotated in Ensembl and counted with featureCounts (version 2.0.1)[10]. All further analyses were performed using R (version 4.1.2). For gene level count data, the R package EDASeq was used to account for sequencing depth (upper quartile normalization)[11]. Latent sources of variation in expression levels were assessed and accounted for using RUVSeq (RUVs) [12]. Appropriate choice of the RUVSeq parameter k was determined through inspection of RLE plots and PCA plots. Differential expression analysis was conducted using edgeR, applying FDR < 0.05 [13]. Statistical significance of the overlap between two groups of genes was performed using the online analysis tool (http://nemates.org/MA/progs/overlap_stats.html). Briefly, two groups of genes are compared and found to have x genes in common. A representation factor and the probability of finding an overlap of x genes are calculated based on the exact hypergeometric probability and normal approximation. Linear regression was applied using Pearson (R) Calculator (https://www.socscistatistics.com/pvalues/pearsondistribution.aspx). Analysis code available through Github at <https://github.com/YannVRB/16p11.2-del-and-3gKO-bulk-RNA-seq.git>

Enrichment analysis of DEG-associated pathways was performed with the Kyoto Encyclopedia of Genes and Genomes (KEGG) database. The analyses were done with the Cytoscape (version.3.9.0, https://cytoscape.org/) plug-in ClueGO (version 2.5.6) [14, 15]. Only the pathways with a p-value<0.05 and gene counts ≥3 were considered as significant and displayed. To connect the terms in the network, ClueGO utilizes kappa statistics in which here was set as ≥0.4. In addition, an overrepresentation test (Fisher’s test corrected for False Discovery Rate) was performed in PANTHER using the reactome pathway annotation dataset (Reactome version 65 Released 2021-10-01). Only pathways with FDR < 0.05 were displayed. Finally, protein-protein interaction (PPI) networks were build using STRING database (version 11.5) with a minimum required interaction score of 0.400. Edges of the network indicated the strength of data support. For better understanding of interactions, the k-mean cluster option in the string database was applied to divide the interacting network into 3 clusters composed of closely connected proteins in the network.

Acquisition and analysis of diffusion-weighed MRI data sets

*Image acquisition*

Male and female mice at an age of PD 42-47 (6-week-old), 70-day-old (10-week-old), or 114-118-day-old (16-week-old) were euthanized using CO2 and perfused using 4% PFA with added 2mM concentration of gadolinium agent and 1μl/ml heparin. Imaging was performed ex vivo using a GE/Agilent Discovery 901 7-Telsa pre-clinical scanner. MRI imaging acquisition consisted of a 32-direction diffusion tensor imaging (DTI) scan to investigate microstructural integrity of white matter fiber tracts. The diffusion-weighed scan utilized an 8-shot 2D segmented echo-planar imaging (EPI) to a 128 × 128 matrix over a 18 mm × 18 mm field of view with 49 slices at 0.3 mm thickness, resulting in a voxel resolution of 0.1406mm x 0.1406mm x 0.3mm. The scan acquired 15 diffusion directions with b = 1000 s/mm² along with 2 T2 (b = 0) images with TR/TE = 8000ms/17.0ms.

*Image preprocessing and TBSS analysis*

After the acquisition, diffusion-weighed images were converted from DICOM to NIFTI format using DCM2NIIX and examined for quality. Bias field correction was applied using N4BiasFieldCorrection from Advanced Normalization Tools (ANTs), images were brain-extracted using hand-drawn masks, and were resampled into an isometric voxel resolution of 0.2 mm3 for ease of use in the processing pipeline. Using the diffusion toolbox (FDT) a tensor model was fit to the diffusion data to generate the fractional anisotropy (FA) images. Voxel-wise statistical analysis was performed on the FA data using a version of Tract-Based Spatial Statistics (TBSS) from FSL that was modified in house and optimized for use with mouse imaging analysis. FA data for all animals were then aligned into Waxholm space19 using the nonlinear registration tool FNIRT from FSL. Next, mean FA images were created and thresholded at a value of 0.2 to delineate major fiber tracts and create a mean FA skeleton representative of the centers of all fiber tracts in the data. Each subject's aligned FA data was projected onto this mean FA skeleton and the resulting data was used to perform voxel-wise cross-subject statistics using the randomize tool from FSL, with 500 permutations and threshold-free cluster enhancement (TFCE) at p < 0.05 to check for significance in the contrasts.

SUPPLEMENTARY FIGURE LEGENDS

Figure S1. Gene modification design. Comparison of DNA sequence between wildtype and mutant *of Taok2* (A), *Sez6l2* (B), and *Mvp* (C).

Figure S2. Nearby genes expression changes of 3g del/+ mice. (A-E) qPCR results demonstrated there is no gene expressions alteration in the mutant mice.

Figure S3. Demographics of 3g del/+ mice inheritance and growth curve. (A) There is no noticeable fatality on 3g del/+ mice. (B) Comparison of developmental milestones of body weight during post-weaning. Male 3g del/+ mice displayed decreased weight (main effect of genotype, F (1, 34) = 4.792, p = 0.0356), but female mice did not (main effect of genotype, F (1, 26) = 0.6839, p = 0.6400). Values are displayed as mean (±SEM) and significance values are set at *p<0.05 and **p<0.01.

Figure S4. Rotarod test on 3g del/+ mice. (A, B) Performance of 3g del/+ mice and wt mice on the accelerating rotarod (4-40rpm) was measured by time to fall off. The latency did not differ among the groups for either males (F (1, 15) = 0.04061, p = 0.8430) or females (F (1, 17) = 0.6865, p = 0.4188).

Figure S5. Body weight during food restriction. (A, B, D, E) After food restricted period, wt and 3g del/+ mice weighed significantly less than ad libitum feeding period (wt male, t(7) = 13.10, p < 0.001; 3g male, t(12) = 23.11, p < 0.001; wt female, t(10) = 11.60, p < 0.001; 3g del/+ female, t(9) = 11.44, p < 0.001). (C, F) The percentage of weight loss in wt and 3g del/+ mice did not differ (male, t(19) = 0.2528, p = 0.8031; female, t(19) = 0.6956, p = 0.4951).

Figure S6. Additional behavioral phenotypes of 3g del/+ mice. (A) male 3g del/+ mice spent significantly more time in the center than wt mice (t(17) = 2.272, p = 0.036), while female 3g del/+ mice did not differ in center activity (t(17) = 0.8626, p = 0.400). The total distance traveled during the task did not differ among the groups for either males or females during open field test (male, t(17) =1.677, p = 0.111; female, t(17) = 1.060, p = 0.304). (B) Both male and female 3g del/+ mice show sniffing preference for the social stimulus versus a novel object (male, main effect of phase, F (1,20) = 59.83, p<0.001; female, main effect of phase, F (1,19) = 30.82, p<0.001). However, genotype difference was not detected in sniffing time with mouse regardless of sex (male, main effect of genotype, F (1, 20) = 2.200, p = 0.154; female, main effect of genotype, F (1, 19) = 0.019, p = 0.892). The total distance traveled during the social approach task did not differ among the groups (male, t(20) =0.597, p = 0.558; female, t(19) = 0.833, p = 0.4152).

Figure S7. Quadrant plot of 248 DEGs with labelled ribosomal genes regulated in 3g del/+ male mice and 16p11.2 del/+ male mice.

Figure S8. Pathway analysis of RNAseq results on 3g del/+ male mice specific DEGs (569 DEGs). (A) KEGG enrichment pathway analysis. (B) PPI networks using STRING database. (C) PANTHER using the reactome pathway annotation dataset.

Figure S9. Pathway analysis of RNAseq results on 16p11.2 del/+ male mice specific DEGs (633 DEGs). (A) KEGG enrichment pathway analysis. (B) PPI networks using STRING database. (C) PANTHER using the reactome pathway annotation dataset.

Figure S10. Pathway analysis of RNAseq results on 3g del/+ female mice specific DEGs (340 DEGs). (A) KEGG enrichment pathway analysis. (B) PPI networks using STRING database. (C) PANTHER using the reactome pathway annotation dataset.

Figure S11. Pathway analysis of RNAseq results on 16p11.2 del/+ female mice specific DEGs (121 DEGs). (A) KEGG enrichment pathway analysis. (B) PPI networks using STRING database. (C) PANTHER using the reactome pathway annotation dataset.

Figure S12. Fiber tract changes in 3g del/+ mice compared to wild types. Increased FA in mutant mice compared to wt mice is represented in blue and decreased FA is displayed in red. (A-D) FA changes are highlighted throughout the brain in 3g del/+ or 16p11.2 del/+ mice (6-week-old) compared to wt mice. (E) FA change is not detected in male 10-week-old 3g mice. (F) Decreased FA changes are detected in male 10-week-old 16p11.2 del/+ mice.

Figure S13. Graphic abstract.

SUPPLEMENTARY TABLE

Table S1. The list of 817 DEGs from 3g del/+ males vs wt males in the striatum.

Table S2. The list of 346 DEGs from 3g del/+ females vs wt females in the striatum.

Table S3. The list of 881 DEGs from 16p11.2 del/+ males vs wt males in the striatum.

Table S4. The list of 127 DEGs from 16p11.2 del/+ females vs wt females in the striatum.

Table S5. The list of 36 overlapping DEGs in the striatum of male and female 3g del/+ mice.

Table S6. The list of 42 overlapping DEGs in the striatum of male and female 16p11.2 del/+ mice.

Table S7. The list of 248 overlapping DEGs in the striatum of male 3g del/+ mice and male 16p11.2 del/+ mice.

Table S8. The list of 6 overlapping DEGs in the striatum of female 3g del/+ mice and female 16p11.2 del/+ mice.
